# NetAct: a computational platform to construct core transcription factor regulatory networks using gene activity

**DOI:** 10.1101/2022.05.06.487898

**Authors:** Kenong Su, Ataur Katebi, Vivek Kohar, Benjamin Clauss, Danya Gordin, Zhaohui S. Qin, R. Krishna M. Karuturi, Sheng Li, Mingyang Lu

## Abstract

A major question in systems biology is how to identify the core gene regulatory circuit that governs the decision-making of a biological process. Here, we develop a computational platform, named NetAct, for constructing core transcription-factor regulatory networks using both transcriptomics data and literature-based transcription factor-target databases. NetAct robustly infers regulators’ activity using target expression, constructs networks based on transcriptional activity, and integrates mathematical modeling for validation. Our in-silico benchmark test shows that NetAct outperforms existing algorithms in inferring transcriptional activity and gene networks. We illustrate the application of NetAct to model networks driving TGF-β induced epithelial-mesenchymal transition and macrophage polarization.

## Background

One of the major goals of systems biology is to infer and model complex gene regulatory networks (GRNs) which underpin the biological processes of human disease^1–6^. Particularly important are those gene networks that control decisions regarding cellular state transitions (*e*.*g*., replicative to quiescent^7–9^, epithelial to mesenchymal (EMT)^10^, pluripotent to differentiated^11,12^), given the central importance of such regulatory processes to both healthy development as well as diseases such as cancer.

To construct and model GRNs associated with the biological process under investigation, researchers have developed two primary systems biology approaches. The first is a *bottom-up approach*, in which researchers focus on identifying a core GRN composed of a small set of master regulators^13^. Once the core GRN is obtained, mathematical modeling is then applied to simulate the gene expression dynamics^14–17^, which helps elucidate the potential gene regulatory mechanism driving the biological process in question. The current practice for synthesizing a core GRN is by compiling data via an extensive literature search, *e*.*g*., in these studies^18–20^. While this works well for systems where sufficient knowledge has been gained and accumulated, it is less effective in cases where key component genes and regulatory interactions have yet to be discovered. Due to rapid increase of biomedical publications, manual synthesis of literature information has become extremely time-consuming and prone to human error in data interpretation. One way to address the labor-intensive issue is to rely on existing manually curated databases, such as KEGG^21^ and Ingenuity Pathway Analysis (IPA)^22^. However, these databases often compile gene regulatory interactions from different tissues, species, or diseases. Therefore, it is hard to obtain context-specific interactions directly from these types of databases.

The second approach adopts a *top-down* perspective, in which researchers apply bioinformatics and statistical methods on genome-wide transcriptomics and/or genomics data to infer large-scale GRNs^13^. These data-driven methods are ideal for obtaining a global picture of gene regulation and the overall structure of gene-gene interactions. This approach also helps to characterize key regulators and regulatory interactions between genes that are specific to the biological context of the study. However, conventional bioinformatics methods for gene network inference are usually not designed to identify an integrated working system. These methods typically rely on significance tests to determine the nodes and edges of a gene network, yet it is rare to evaluate whether the constructed gene network is capable of operating as a functional dynamical system^23^. Moreover, many statistical methods work well to identify the association between genes, but not their causation, thus limiting the applicative value of the top-down approach in characterizing gene regulatory mechanisms.

To overcome the above-mentioned issues, a relatively new approach has been explored in several studies in which the top-down and bottom-up approaches are integrated to infer and model a core GRN^23– 31^. In this combined approach, a GRN is constructed with bioinformatics tools using genome-wide gene expression data, followed by mathematical modeling of the GRN to simulate gene expression steady states and explore their similarity with biological cellular states. The simulations can help validate the accuracy of the constructed GRN and further clarify the regulatory roles of genes and interactions in driving cellular state transitions. This combined approach helps to discover existing and new regulatory interactions specific to the cell types and experimental conditions under study. Additionally, it helps pinpoint master regulators and reduce the system’s overall complexity. The GRN modeling is particularly crucial for cases with non-trivial cellular state transitions, such as multi-step state transitions as observed in Epithelial-Mesenchymal Transition (EMT)^32^, and bifurcating state transitions, as observed in stem cell differentiation^33^. This is because the GRNs constructed by the top-down approach are not guaranteed to capture these state transition patterns. So far, to the best of our knowledge, there is no computational platform available that utilizes this combined approach for systematic GRN inference and modeling.

In this study, we introduce a computational platform, named NetAct, for inferring a core GRN of key transcription factors (TFs) using both transcriptomics data and a literature-based TF-target database. Integrating both resources allows us to take full advantage of the existing knowledgebase of transcriptional regulation. NetAct adopts the combined top-down bioinformatics and bottom-up systems biology approaches, designed specifically to address the following two major issues.

First, many network inference methods rely on correlations of gene expression data, yet the actual transcriptional activities of many master regulators may not be reflected in their gene expression. Instead, the activity may be better associated with either their protein level, the level of a certain posttranslational modification, localization, or their DNA binding affinity. As a result, the master regulators with weak correlations between the expression level and the transcriptional activity will likely be discarded in the network. Some algorithms have been developed to infer the activities of regulators from transcriptomics data, such as VIPER^34^, NCA^35^, AUCELL^36^. However, most of these algorithms 1) are not designed for gene network modeling, or 2) still rely on coexpression of a TF and its targeted genes, or 3) do not take advantage of known regulatory interactions from the literature, hindering their applicability as automated algorithms for generic use in systems biology.

Second, conventional mathematical modeling approaches have been applied over the years to simulate the dynamics of a GRN, yet they are not particularly effective in analyzing core GRNs. A popular method models the gene expression dynamics of a system using the chemical rate equations that govern the associated gene regulatory processes. However, it is difficult to directly measure most of the kinetic parameters of a GRN. Although some parameter values can be learned from published results, many others are often based on educated guesses which significantly limits the predictive power of mathematical modeling. Moreover, a core GRN is not an isolated system. Thus, an ideal modeling paradigm should also consider other genes that interact with the core network. To address this infamous parameter issue, we have developed the modeling algorithm RACIPE^29,37,38^ in previous work that analyzes a large ensemble of mathematical models with random kinetic parameters. RACIPE has been applied to model the dynamical behavior of gene regulatory networks of different biological processes, such as epithelial-mesenchymal transition^23,29^, cell cycle^38^, stem cell differentiation^39^.

The new NetAct platform addresses the above-mentioned issues by (1) inferring the activities of TFs for individual samples using the gene expression levels of their targeted genes, (2) identifying the regulatory interactions between two TFs based on their activities rather than their expressions, (3) and subsequently simulating the constructed core GRN with RACIPE to validate and evaluate the gene expression dynamics of the core GRN. In this paper, we describe in detail the NetAct platform, extensive benchmark tests for TF-target databases, TF activity inference, and network construction, and two examples of applications to model GRNs with time series gene expression data.

## Results

We developed a computational systems-biology platform, named NetAct, to construct transcription factor (TF)-based GRNs using TF activity. The method uniquely integrates both generic TF-target relationships from literature-based databases and context-specific gene expression data. NetAct also integrates our previously developed mathematical modeling algorithm RACIPE to evaluate whether the constructed network functions properly as a dynamical system. It evaluates the roles of every gene in the network by in-silico perturbation analysis. NetAct has three major steps: (1) identifying the core TFs using gene set enrichment analysis (GSEA)^40^ with an optimized TF-target gene set database (Fig. 1a); (2) inferring TF activity (Fig. 1b); (3) constructing a core TF network (Fig. 1c). Then, the network is validated and analyzed by simulating its dynamics using mathematical modeling by RACIPE (see Supplemental Material SI5). Details of each step is given in the Methods section and Supplemental Material. Below, we demonstrate how we optimized the NetAct algorithm, compared its performance of activity inference with three existing methods using in-silico gene expression data, and applied the network modeling approach to two biological datasets.

**Fig.1.**
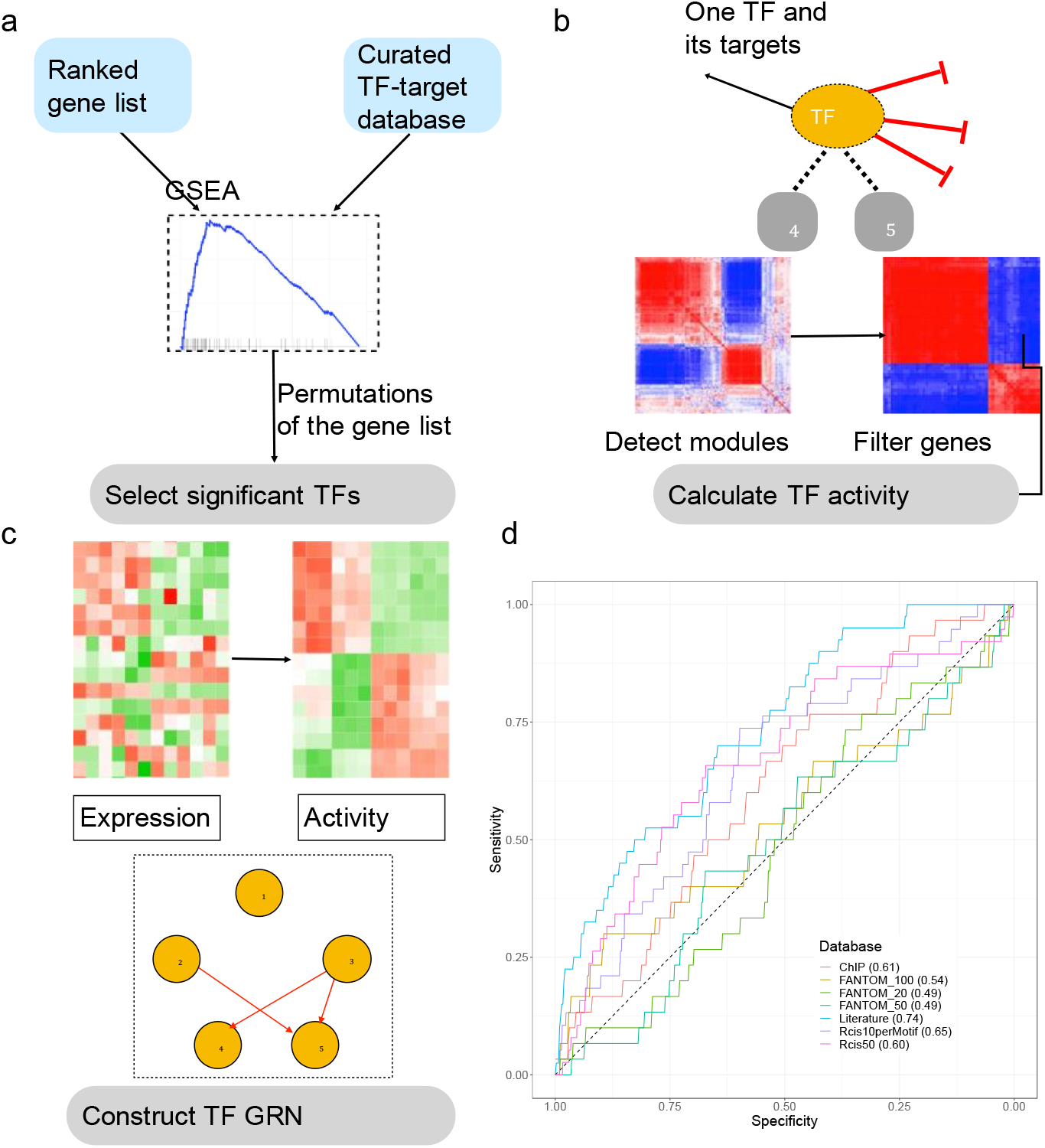
Schematics of NetAct. (**a**) First, key transcription factors (TFs) are identified using gene set enrichment analysis (GSEA) with a literature-based TF-target database. (**b**) Second, the TF activity of an individual sample is inferred from the expression of target genes. From the co-expression and modularity analysis of target genes, we find target genes that are either activated (blue), inhibited (red), or not strongly related to the TF (grey). The activity is defined as the weighted average of target genes activated by the TF minus the weighted average of target genes inhibited by the TF. (**c**) Lastly, a TF regulatory network is constructed according to the mutual information of inferred TF activity and literature-based regulatory interactions. (**d**) Performance of GSEA for various TF-target gene set databases. The plot shows the sensitivity and specificity with different q-value cutoffs. The gene set databases in the benchmark include the combined literature-based database (D1), FANTOM5-based databases (D2) with 20, 50, 100 target genes per TF, the combined experimental-based database (D3, ChIP), and RcisTarget databases (D4), one with 10 targets per TF binding motif and another with 50 total number of targets per TF.

### Literature-based TF-target relationships facilitate TF inference

To establish a comprehensive gene set database containing TF-target relationships, we considered data from different sources (Table S1, also see Supplemental Material SI1). They are (D1) a literature-based database, consisting of data from TRRUST^41^, RegNetwork^42^, TFactS^43^, and TRED^44^; (D2) a gene regulatory network database FANTOM5^45^, whose interactions are extracted from networks constructed using RNA expression data from 394 individual tissues; (D3) a database derived from resources of putative TF binding targets, including ChEA^46^, TRANSFAC^47^, JASPAR^48^, and ENCODE^49^; and (D4) a database derived from motif-enrichment analysis, RcisTarget^50^. These databases have been frequently used to study transcriptional regulations and have already been utilized for network construction^29,51^.

We evaluated the performance of these databases by GSEA on a benchmark dataset. GSEA is a popular statistical method that can be used to evaluate significant overlapping between a set of genes and differentially expressed genes between two experimental conditions. Using various types of TF-target databases, our goal is to find the best version of the database, so that GSEA can detect the target gene sets of the relevant TFs to be statistically significant. This benchmark dataset, denoted as *set B*, consists of a compilation of 11 microarray and 27 RNA-Seq gene expression data (Table S2). Each of these datasets contains at least three samples under the normal condition (control) and three samples under the treatment condition in which a specific TF is treated by knockdown (KD). We applied GSEA (with slight modifications, details in Methods) on the set *B* to evaluate whether the enrichment analysis can detect the perturbed TFs. The underlying assumption is that, with a better TF-target gene set database, GSEA will be more likely to detect the corresponding perturbed TFs. For each TF-target database and each gene expression data in set *B*, we calculated the q-values of all the TFs in the database by GSEA to determine whether the target genes of the perturbed TF are enriched in the differentially expressed genes. We found that more significant q-values are usually associated with relatively larger number of targets for each TF; however, too many (*e*.*g*., greater than 2000) targets will result in non-significant q-values. The summary statistics, such as the total number of TFs and the average number of target genes per TF, are summarized in Table S1. Furthermore, these corresponding q-values from all the gene expression data are converted to specificity and sensitivity values (see Methods), and different databases are compared based on the area under the sensitivity-specificity curves (Fig. 1d). We found that the literature-based database has the best overall performance, thus we used this database for further analyses. Our results are in line with a previous benchmark study^52^ that literature-based TF-target database outperforms others in capturing transcriptional regulation.

### Inferring TF activity without using TF expression

NetAct can accurately infer TF activity for an individual sample directly from the expression of genes targeted by the TF (see Methods). In the following, we will illustrate how NetAct infers TF activity on two cases of microarray KD experiments -- one case for shRNA KD of FOXM1 and shRNA KD of MYB in lymphoma cells (GEO: GSE17172^53^), and another case for KD of BCL6 on both OCI-Ly7 and Pfeiffer GCB-DLBCL cell lines (GEO: GSE45838^34^). NetAct first successfully identified the TFs that undergo knockdown in each case, *i*.*e*., FOXM1, MYB and BCL6 respectively, by applying GSEA on the optimized TF-target database (q value < 0.15).

Next, for each identified TF, NetAct calculates its activity using the mRNA expression of the direct targets of the TF. We first constructed a Spearman correlation matrix from the expression of the targeted genes. As shown in Fig. 2a, the correlation matrix after hierarchical clustering analysis typically consists of two red diagonal blocks, two blue off-diagonal blocks, and the remaining elements with low correlations which will be filtered out subsequently (details in Methods). Within the red blocks, the expression of any column gene is positively correlated with that of any row gene; while within the blue blocks, the expression of any column gene is negatively correlated with that of any row gene. This indicates that the genes in the two red blocks are anti-correlated in gene expression with each other. However, if the correlation matrix is constructed from 100 or 200 randomly selected genes (Fig. 2bc), such a clear pattern disappears. Thus, our observation suggests that genes from one of the red blocks are activated by the TF, whereas genes from the other block are inhibited by the TF. Moreover, filtered genes are not likely to be directly targeted by the TF in this context, or they are regulated by multiple factors simultaneously and are thus likely not a good indicator for the TF activity.

**Fig.2.**
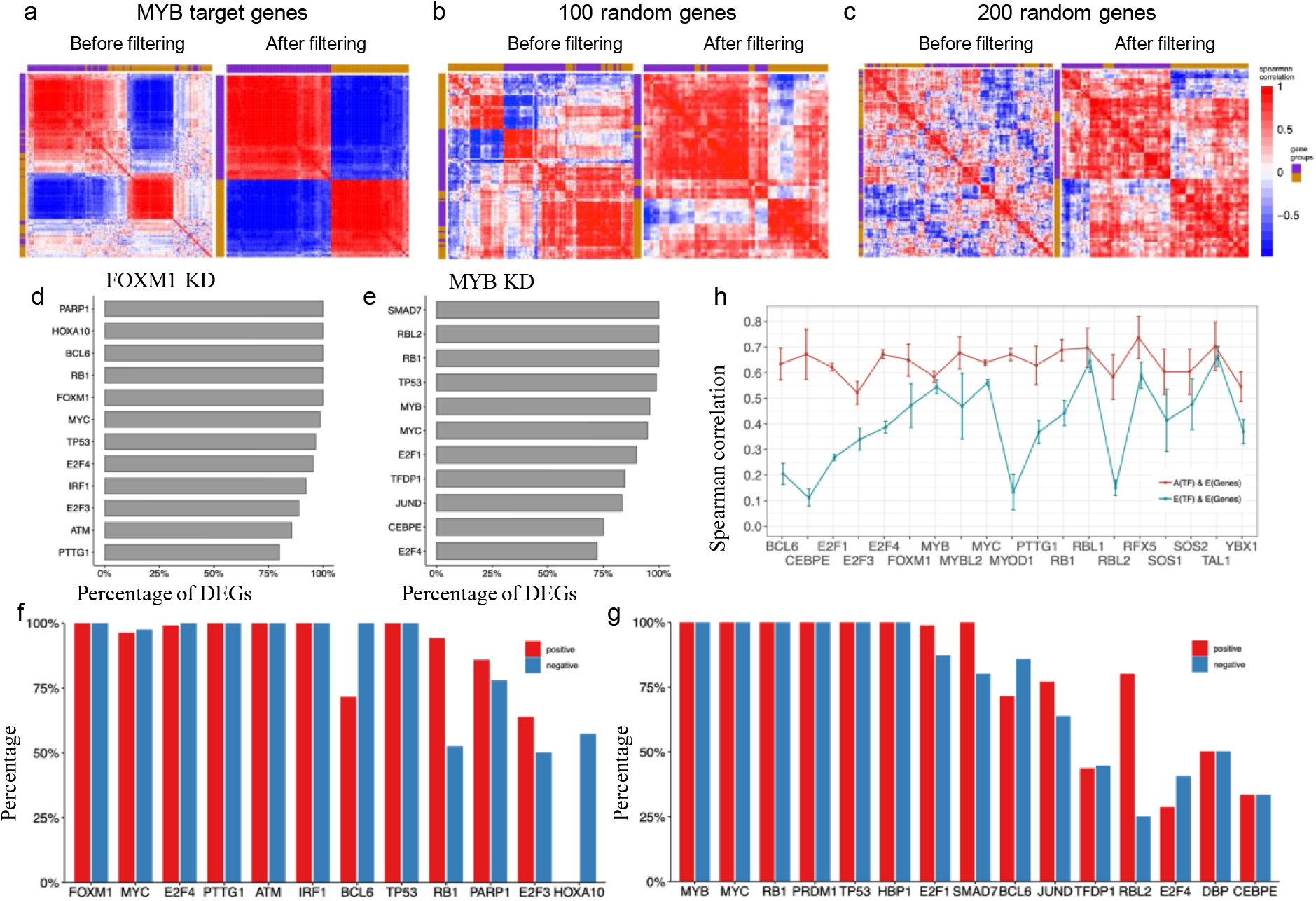
Illustration of the grouping scheme for target genes of a transcription factor. (**a**) shows the co-expression matrix of MYB target genes in shRNA knockdown of MYB lymphoma cells by hierarchical clustering analysis (Pearson correlation and complete linkage). (**b, c**) demonstrate the poor clustering results from the co-expression of randomly selected 100 (in **b**) and 200 genes (in **c**). In panels **(a – c)**, the left subplots show the outcomes of all tested genes, and the right subplots show the outcomes of genes after the filtering step. Compared to the random cases, MYB target genes have a clear pattern of red and blue diagonal blocks from their co-expression. (**d, e**) show the percentage of differentially expressed genes remained after the filtering step in the case of FOXM1 and MYB knockdown, respectively. (**f, g**) show the proportion of genes from the activation group that are positively correlated with the TF expression (red bars) and the proportion of genes from the inhibition group that are negatively correlated with the TF expression (blue bars). (**h**) Pearson correlation (average and standard deviation) between TF activity and target expression (red) and between TF expression and target expression (blue).

We further evaluated how the filtering step removes noise and retains the important genes in the analysis. We found that, after the filtering step, most of the differentially expressed (DE) genes are retained, as evidenced by Fig. 2d. Here, DE genes from each comparison were retrieved by using *limma* with a cutoff for the adjusted p-values at 0.05 and a cutoff for the log2 fold changes at 2. Subsequently, for DE TFs we evaluated the Spearman correlations between the TFs and the corresponding targeted genes. In traditional approaches (such as ARACNe^1^, WGCNA^54^, and BEST^55^), the co-expression between a TF and its targeted genes are commonly used to identify its association and assign the sign (activation or inhibition) of the regulation. We found that, for each TF, most of the genes in a block either positively correlate with the TF expression (Fig. 2fg, blue bars), or they negatively correlate with the TF expression (Fig. 2fg, red bars). The tests demonstrate that, without directly using TF expression, NetAct can successfully identify two groups of important target genes – genes in each group are either activated or inhibited by the TF. These two groups of genes are further used to infer TF activity by a weighted average of their gene expression (Equation 1 in Methods). Additionally, we found that the correlations between inferred TF activity and target expression are usually higher than the correlations between TF expression and target expression (Fig. 2h).

### Evaluating activity inference and network construction in a simulation benchmark

To evaluate the accuracy and robustness of inferred TF activity, we performed extensive benchmark tests to compare NetAct with other existing methods. We first performed the benchmark tests on simulated data because TF activity is usually not directly measurable. The activity of a TF can be related to its protein level or the level of a particular posttranslational modification, such as phosphorylation. Therefore, it is very difficult to obtain the ground truth of TF activity from an experimental data set. Thus, in this benchmark test, we rely on mathematical modeling to simulate both the expression and activity of each TF from a synthetic TF-target network. With this simulated data, we benchmark NetAct against other methods.

To establish the simulated benchmark data set, we first constructed a synthetic TF-target network with a total of 30 TFs. Each TF has 20 target genes randomly selected with replacement from a pool of 1000 genes. In addition, each TF also regulates two (randomly selected) of the 30 TFs. This synthetic network has a hierarchical structure, where a target gene may be co-regulated by multiple TFs. The type of each TF-to-TF regulation is either excitatory, inhibitory, or signaling, with a chance of 25%, 25%, and 50%, respectively; the type of each TF-to-target regulation is either excitatory or inhibitory with a 50% chance for each. Here, the signaling regulation changes the activity of a TF without changing its expression; whereas the excitatory or inhibitory interactions changes both of the activity and expression. From one realization of the synthetic network generation, the final synthetic network contains a total of 477 genes (30 TFs, 447 targeted genes) and 660 regulatory links (Fig. 3a). See Supplemental Material SI4 for more details.

**Fig. 3.**
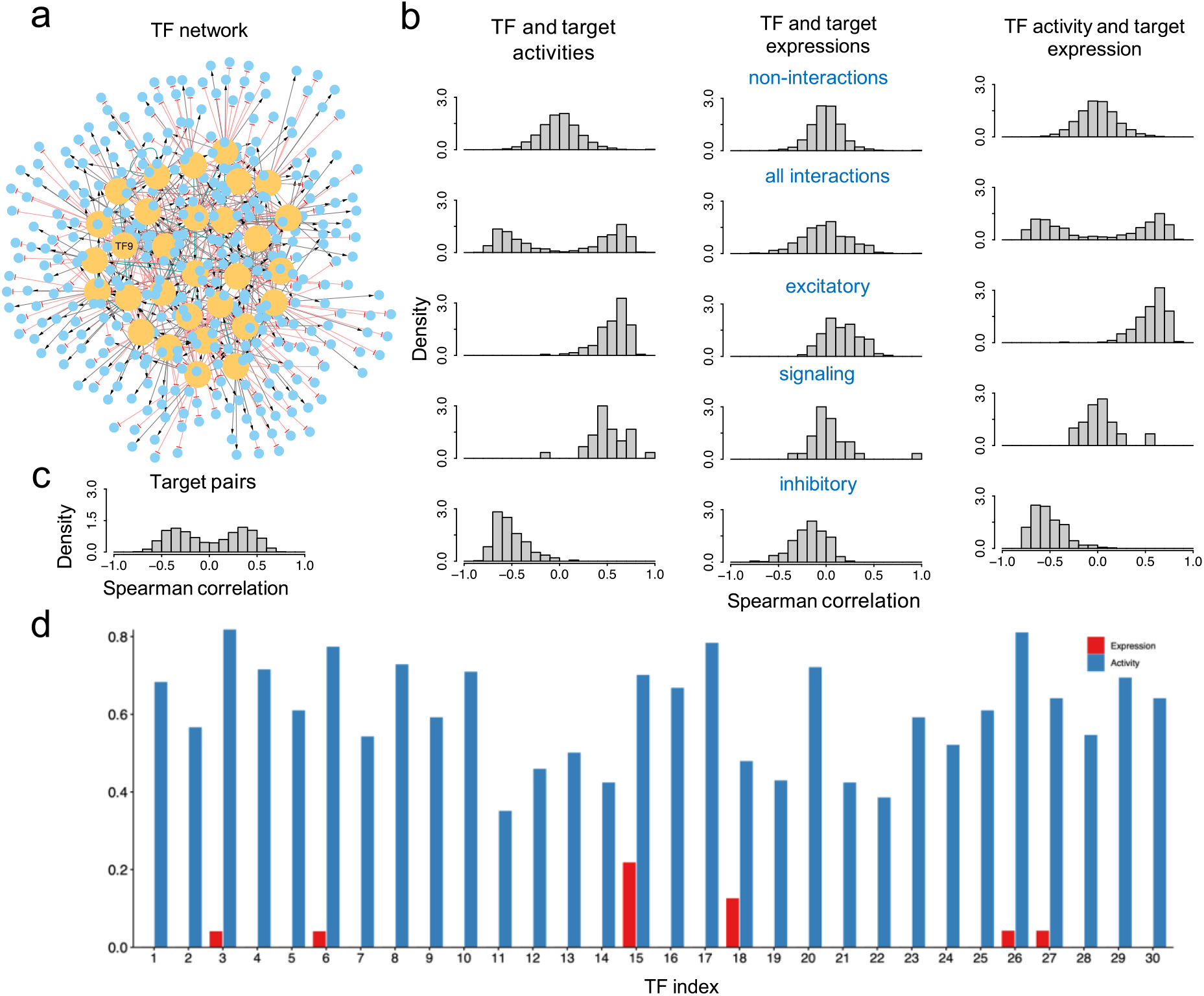
Simulation of both gene expression and activity of a synthetic GRN. (**a**) shows the synthetic GRN consisting of 30 TFs and 447 target genes. An edge of transcriptional activation is shown as black line with an arrowhead; an edge of transcriptional inhibition as red line with a blunt head; an edge of signaling interaction as green line with an arrowhead. Transcription factor labeled as TF9 was selected for knockdown simulations. (**b**) shows the summary of the correlation analyses of the simulated expression and activity. The left, middle, and right columns represent the outcomes for TF and target activities, TF and target expressions, and TF activities and target expressions, respectively. For each category, the histograms of Spearman correlations are shown for non-interacting gene pairs (first row), interacting gene pairs (second row), gene pairs of excitatory transcriptional regulation (third row), gene pairs of excitatory signaling regulation (fourth row), gene pairs of inhibitory transcriptional regulation (fifth row). Here, the target activity is set to be the same as the target expression for non-TF genes. (**c**) shows the histograms of Spearman correlations for gene pairs of target genes from the same TF. (**d**) Jaccard indices between the ground-truth regulons of the synthetic GRN and the regulons inferred by ARACNe using either the simulated expression (red) or activity data (blue).

To simulate the gene expression of the TF-target network, we applied a generalized version of the mathematical modeling algorithm, RACIPE^38^. Using the network topology as the only input, RACIPE can generate an ensemble of random models, each corresponds to a set of randomly sampled parameters. Here, we used RACIPE to generate simulated data including gene expression and TF activity for benchmark. Some previous studies have also adopted a similar modeling approach for benchmarking^56,57^. To consider the effects of a signaling regulatory link, we generalized RACIPE to simulate both expression and activity for each TF. See Supplemental Material SI5 for more details.

In the benchmark test, we used RACIPE to simulate 100 models with randomly generated kinetic parameters. From these 100 models we obtained 83 stable steady-state gene expression and activity profiles for the 477 genes. As expected, TF activity and target activity from a regulatory link are correlated (1^st^ column, 2^nd^ row in Fig. 3b); TF activity and target expression (3^rd^ column, 2^nd^ row in Fig. 3b) are correlated; and the expression of two target genes (Fig. 3c) are correlated. However, there is no strong correlation between TF expression and target expression (2^nd^ column, 2^nd^ row in Fig. 3b) and, for a signaling regulatory link, between TF activity and target expression (3^rd^ column, 4^th^ row in Fig. 3b). Next, we applied ARACNe to predict the regulon (*i*.*e*., the list of targeted genes by a specific TF) using either the simulated expression profiles or the simulated activity profiles. We found that the regulons predicted from the activity profiles are substantially more similar to the predefined regulons (measured by the Jaccard similarity^58^) than those predicted from the expression profiles (Fig. 3d). The results indicate the need of using the TF activity, instead of TF expression, to identify TF-target relationships.

Next, we compared the performance of NetAct with several related algorithms, NCA, VIPER, and AUCell, in inferring TF activity using both the simulated expression profiles from the 83 models and a predefined regulon (*i*.*e*., the association of each TF with its target genes) (details for the implementation of these algorithms in Supplemental Information SI3). The predicted activity was then compared with the simulated activity (ground truth) to evaluate the performance. To mimic the real-life scenario where the target information may not be complete and accurate, we consider more challenging tests where the regulon data is randomly perturbed. Here, for a specific perturbation level, we generated 100 sets of regulon data by replacing a certain number of target genes for each TF with non-interacting genes. The numbers of replaced genes are 0 (0% level of perturbation), 5 (25%), 10 (50%) and 15 (75%), respectively, in different tests. We then evaluated the performance of NetAct, NCA, and VIPER. AUCell protocol advises to include the target genes with only positive interactions in the regulons. To satisfy this criterion, we updated the regulons for both unperturbed and perturbed regulons. For the unperturbed regulons, we retained only the positive interactions; for the perturbed regulons, we retained the positive target genes that were not replaced and a random half of the replaced target genes (assuming that half of the genes are positively regulated by the TF). We then evaluated AUCell performance using these updated regulons (denoted AUCell 1) and non-updated regulons (denoted AUCell 2). As shown in Fig. 4a (also Figs. S3-S6), NetAct significantly outperforms each of the other methods in reproducing the simulated activity profiles at each perturbation level. As expected, the performance of NetAct is decreased by increasing the perturbation levels of the regulon data; however, NetAct still performs reasonably well even when only 25% of the actual target genes are kept in the regulon data. The results indicate that NetAct can robustly and accurately infer TF activity even with a noisy TF-target database.

**Fig. 4.**
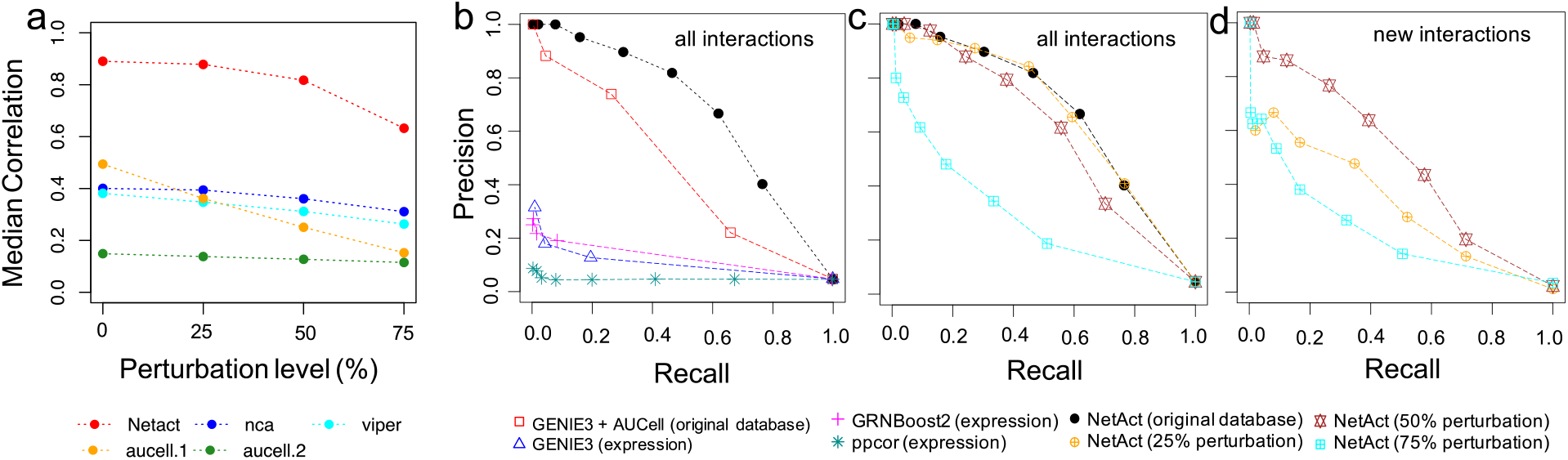
The performance of activity and network inference from a simulation benchmark. **(a)** TF activity inference. TF activity was inferred by several methods using the gene expression data simulated from the synthetic TF-target gene regulatory network (GRN) and the corresponding regulons. For each TF, we computed Spearman correlations between the inferred activity and simulated activity (ground truth) for all the simulated models. Then, we calculated the average correlation values over all TFs. The plots show the median of average correlations for the cases where we used the original regulons defined by the TF-target network (0% perturbation), and the regulons where 5 (25% perturbation), 10 (50% perturbation), and 15 (75% perturbation) target genes are randomly replaced with non-interacting genes, respectively. The median values were computed over 100 repeats of random replacement for each perturbation level, and the values of the average correlations are reported for the case of zero perturbation. Shown are the results for NetAct (red), NCA (blue), VIPER (cyan), AUCELL 1 where regulons contain only positively associated target genes (orange), and AUCELL 2 where regulons contain all target genes (green). **(b-d)** Network inference. The panels show the performance of network inference algorithms from the simulation benchmark by the precision and recall for different link selection thresholds. **(b)** Network inference performance against all ground-truth regulatory interactions. Tested methods are GENIE3, GRNBoost2, and PPCOR, using transcription factor (TF) expression; GENIE3 using TF activity inferred by AUCell; NetAct using its inferred TF activity. For the latter two methods, original (unperturbed) regulons obtained from the regulatory network were used. **(c)** Network inference performance of NetAct against all ground-truth regulatory interactions using the regulons with 0% (the original), 25%, 50%, and 75% target perturbations. **(d)** Network inference performance of NetAct in discovering new regulatory interactions not existing in the regulons. NetAct was applied using the regulons at different perturbation levels (25%, 50%, and 75%). The benchmark results shown here are for the case of the untreated simulation. The results for the case of the knockdown simulation are shown in **Fig. S7**.

Furthermore, we tested another scenario where the test data contains simulated data from two experimental conditions, *e*.*g*., one representing an unperturbed condition and the other representing a perturbed condition. Here, we used the same synthetic network but compiled 40 expression and activity data from the above-mentioned simulation (unperturbed condition), together with 43 expression and activity data from the simulations in which a specific TF (TF9) is knocked down (perturbed condition). We then performed a similar test as above and found that NetAct outperformed each of the other methods (Fig. S2 and Fig. S7a). The notable performance gain of NetAct mainly emanates from the removal of incoherent (or noisy) targets of a TF before the activity calculation in NetAct (see Methods).

In addition, we performed a network construction benchmark of NetAct and a few other network construction algorithms using the in-silico simulation data set, as shown in Fig. 4bcd. NetAct, using the TF activity inferred from the original regulon database, outperforms not only network construction methods using gene expression, such as GENIE3^59^, GRNBoost2^60^, and ppcor^61,62^, but also GENIE3 using the TF activity inferred by AUCell (Fig. 4b). The last approach was presented to mimic a popular method SCENIC. Moreover, we evaluated the performance of NetAct when using a perturbed regulon database. We found that NetAct remains performing well when the perturbation level is as large as 50%, when evaluated by all the ground-truth interactions (Fig. 4c) and by those not presented in regulon database (Fig. 4d). The latter case was designed to evaluate the capability of NetAct in predicting novel interactions. We observed similar outcomes for the case of the second scenario of the simulation data from two conditions (Fig. S7bcd) (see Supplemental Information SI6 for details of the benchmark method). In summary, our in-silico benchmark test demonstrates the high performance of NetAct over existing state-of-the-art methods in both inferring TF activity and gene regulatory networks.

### Characterizing cellular state transitions by GRN construction and modeling

In the previous sections, we demonstrated the capability of NetAct in identifying the key TFs and predicting TF activity. With these data, NetAct further constructs a TF-based GRN using the mutual information (MI) of the activity from the identified TFs (details in Methods). We then applied RACIPE to the constructed network to check whether the simulated network dynamics are consistent with experimental observations. In the following, we show the utility of NetAct with two biological examples: epithelial-mechanical transition (EMT) and macrophage polarization.

In the first case (EMT), we analyzed a set of time-series microarray data on A549 epithelial cells undergoing TGF-β induced epithelial-mesenchymal transition (EMT) (GEO: GSE17708)^63^. According to the overall structure of the transcriptomics profiles, we arranged samples from different time points into three groups – early stage (time points 0h, 0.5h and 1h), middle stage (time points 2h, 4h, and 8h) and late stage (time points 16h, 24h, and 72h). We then performed three-way GSEA with our human literature-based TF-target database to identify enriched TFs that are active between either early-middle, early-late and middle-late timepoints. Forty-one TFs (q-value cutoff 0.01) were identified including many major transcriptional master regulators, such as BRCA1, CTNNB1, MYC, TWIST1, TWIST2 and ZEB1, and factors that are directly associated with TGF-β signaling pathway, such as SMAD3^64^, FOS and JUN^65^. The hierarchical clustering analysis (HCA) of the expression and activity profiles for these TFs is shown in Fig. 5a. While the expression profiles are quite noisy, the activities show a clear gradual transition from the epithelial to mesenchymal (M) state. Note that the signs of the activity of a few non-DE TFs were flipped according to experimental evidence of protein-protein interactions and the nature of transcriptional regulation (see Methods for detailed procedures and Table S3 for a list of the changes).

**Fig. 5.**
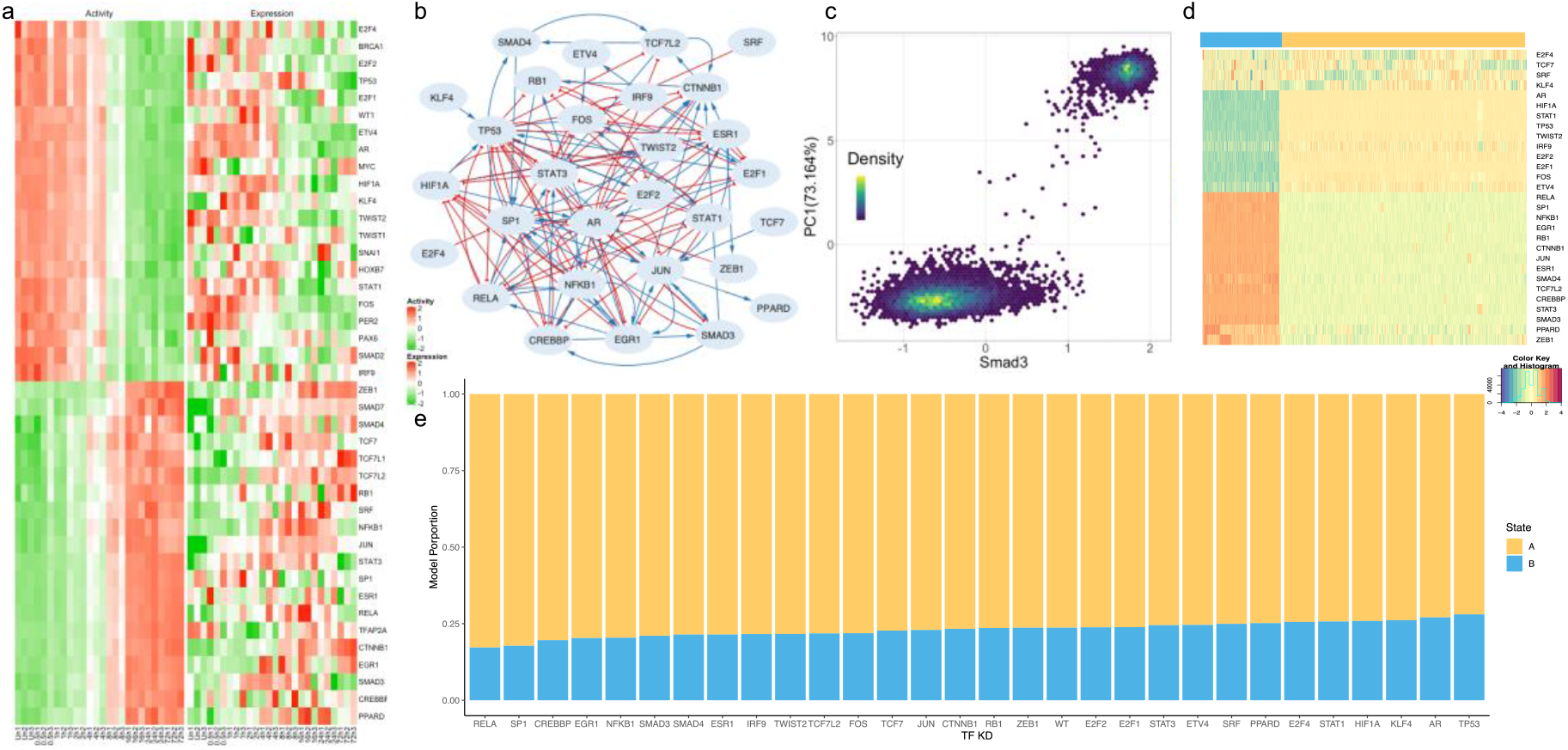
Network modeling of TGF-β induced EMT. Application of NetAct to an EMT in human cell lines using time-series microarray data. (**a**) Experimental expression and activity of enriched transcription factors. **(b)** Inferred TF regulatory network. Blue lines and arrowheads represent gene activation; Red lines and blunt heads represent gene inhibition. **(c)** The relationship between SMAD3 gene activity and the first principal component of the activity of all network genes from RACIPE simulations. **(d)** Hierarchical clustering analysis of simulated gene activity (with Pearson correlation as the distance function and Ward.D2 linkage method). Colors at top indicate the two clusters from the simulated gene activity. The blue cluster represents the mesenchymal state, and the yellow cluster represents the epithelial state. The color legend for the heatmap is at the bottom right. **(e)** Knockdown simulations of the TF regulatory network. The bar plot shows the proportion of RACIPE models in each state (epithelial or mesenchymal) for the conditions of the knockdown of every TF.

We then constructed a TF regulatory network (Fig. 5b) and performed mathematical modeling to simulate the dynamical behavior of the network using RACIPE (Fig. 5cd). We found that, consistent with the expression and activity profiles (Fig.5a), the network clearly allows two distinct transcriptional clusters that can be associated with E (the yellow cluster in Fig.5d) and M states (the blue cluster in Fig.5d). To assess the role of TGF-β signaling in inducing EMT, we performed a global bifurcation analysis^29^ in which the SMAD3 level is used as the control parameter (Fig. 5c). Here, SMAD3 was selected as it is the direct target of TGF-β signaling^64^. As shown in (Fig. 5c), when SMAD3 level is either very low or high, the cells reside in E or M states. However, when SMAD3 is at the intermediate level, the cells could be driven into some rare hybrid phenotypes. These results are consistent with our previous studies on the hybrid states of EMT^32,66^. Using RACIPE, we systematically performed perturbation analyses by knocking down every TF in the network. Our simulation results (Fig. 5e) suggest that knocking down TFs, such as RELA, SP1, EGR1, and CREBBP, *etc*., has major effects in driving M to E transition (MET), while knocking down TFs, such as TP53, AR, and KLF4, *etc*., has major effects in driving E to M transition (EMT). These predictions are all consistent with existing experimental evidence (Table S4).

Compared to a previous model of the EMT network based on an extensive literature survey^19^, the GRN constructed by NetAct identified some of the same regulators induced by the TGF-β pathway, such as SMAD3/4, TWIST2, ZEB1, CTNNB1, NFKB1, RELA, FOS and EGR1. Because of the lack of microRNAs and protein-protein interactions in the database, NetAct didn’t identify factors like miR200 and signaling molecules like PI3K. Interestingly, the NetAct model identifies STAT1/3, which was connected to other signaling pathways, such as HGF, PDGF, IGF1and FGR, but not TGF-β in the previous network model. In addition, the NetAct model identified regulators in other important pathways in TGF-β-induced EMT in cancer cells, *e*.*g*., cell cycle pathway (RB1 and E2F1) and DNA damage pathway (P53).

In the second case, we studied the macrophage polarization program in mouse bone-marrow-derived macrophage cells using time series RNA-seq data (GEO: GSE84517)^67^. In this experiment, macrophage progenitor cells (denoted as UT condition) were treated with (1) IFNγ to induce a transition to the M1 state; (2) IL4 to induce a transition to the M2 state; (3) both IFNγ and IL4 to induce a transition to a hybrid M state. Here, we reprocessed the raw counts of RNA-seq with a standard protocol (details in Supplemental Material SI2). From principal component analysis (PCA) on the whole transcriptomics (Fig.6b), we found that the gene expression undergoes distinct trajectories when macrophage cells were treated with either IFNγ (M1 state) or IL4 (M2 state). When both IFNγ and IL4 were administered, the gene expression trajectories are in the middle of the previous two trajectories, suggesting that cells are in a hybrid state (hybrid M state). We aim to use NetAct to elucidate the crosstalk in transcriptional regulation downstream of cytokine-induced signaling pathways during macrophage polarization.

Here, we applied GSEA on six comparisons – untreated versus IFNγ treated samples (one comparison between the untreated and the treated after two hours, another between the untreated and the treated after four hours, same for the other comparisons), untreated versus IL4 treated samples, and untreated versus IFNγ+IL4 treated samples. Using our mouse literature-based TF-target database, we identified 79 TFs (q-value cutoff 0.05 for UT vs IL4-2h and 0.01 for all others). The expression and activity profiles of these TFs (Fig. 6abc) captures the essential dynamics of transcriptional state transitions during macrophage polarization as follows. NetAct successfully identified important TFs in these processes, including Stat1, the major target of IFNγ, Stat2, Stat6, Cebpb, Nfkb family members, Hif1a and Myc^68–70^. Myc is known to be induced by IL-4 at later phases of M2 activation and required for early phases of M1 activation^69^. Interestingly, we find Myc has high expression in both IL4 stimulation and its co-stimulation with IFN but its activity is high only in IL4 stimulation. We then constructed a TF regulatory network that connects 60 TFs (Fig. 6d) and simulated the network with RACIPE, from which we found that simulated gene expression (Fig. 6f) matches well with experimental gene expression data (Fig.6a) (see Supplemental Information SI7). RACIPE simulations display disparate trajectories from UT to IL4 or IFNγ activation and stimulation with both IL4 and IFNγ. Strikingly, we found in the simulation that there is a spectrum of hybrid M states between M1 and M2 (Fig. 6e), which is consistent with experimental observations of macrophage polarization^68^. Moreover, we also predict from our GRN modeling that the transition from UT to hybrid M is likely to first undergo a transition to either M1 or M2 before a second transition to hybrid M (Fig. 6e). This is because of our observation from the simulation data that there are fewer models connecting UT and hybrid M than any of the other two routes (*i*.*e*., UT to M1, and UT to M2) (Fig. S10). Taken together we showed that the NetAct-constructed GRN model captures the multiple cellular state transitions during macrophage polarization.

**Fig 6.**
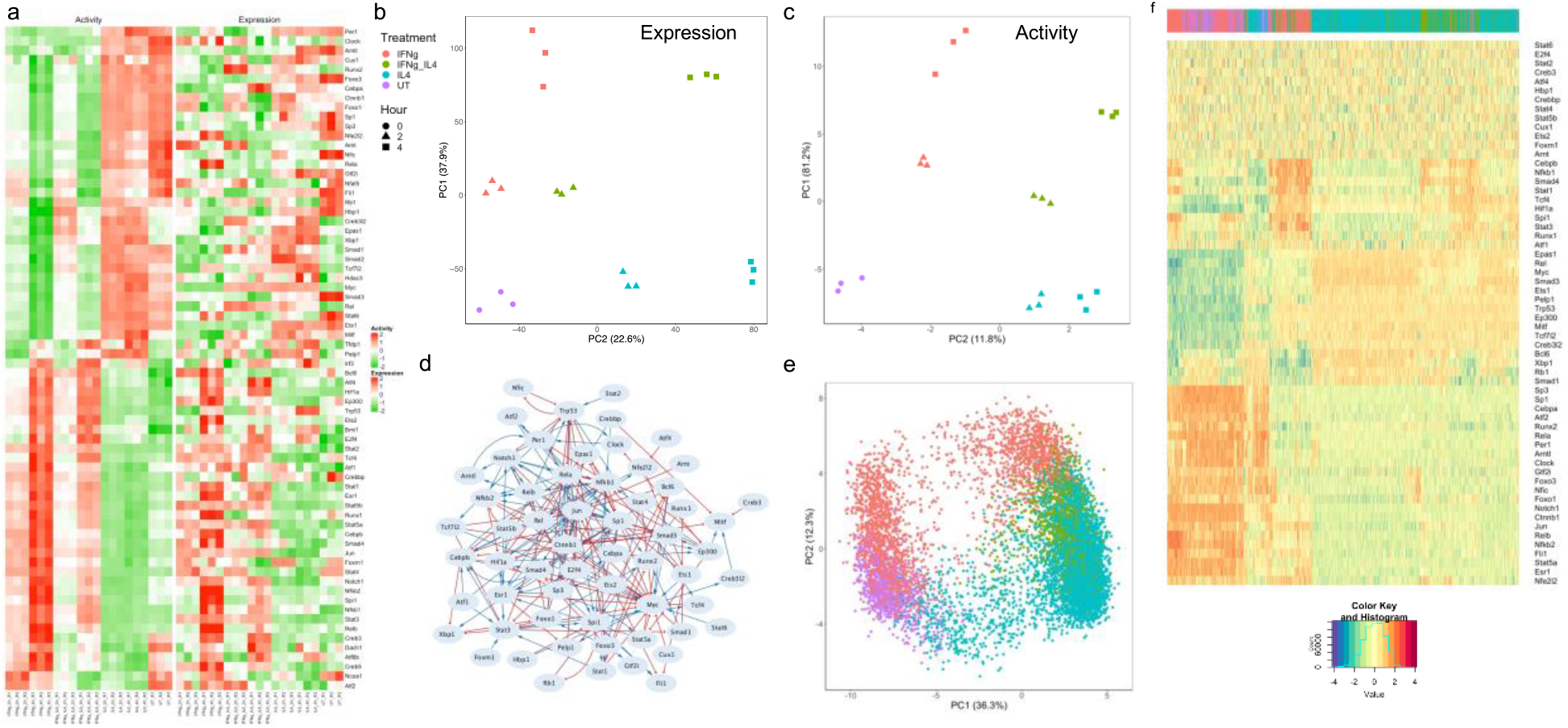
Network modeling of macrophage polarization. Application of NetAct to induced macrophage polarization via drug treatment in mice using RNA-seq data. **(a)** Experimental expression and activity of enriched TFs. **(b)** PCA projection of genome-wide gene expression profiles. Different point shapes indicate time after treatment, and colors indicate treatment types **(c)** PCA projection of gene activity of enriched TFs. **(d)** Inferred TF regulatory network. Blue lines and arrowheads represent gene activation; Red lines and blunt heads represent gene inhibition. **(e)** PCA projection of simulated gene activity of inferred network colored by mapping each model back to experimental data. **(f)** Hierarchical clustering analysis of simulated gene activity (with Pearson correlation as the distance function and Ward.D2 linkage method). Colors at top indicate the mapped experimental conditions. The color legend of the heatmap is at the bottom.

In conclusion, we show that NetAct can identify the core TF-based GRN using both the literature-based TF-target database and the gene expression data. We also demonstrate how RACIPE-based mathematical modeling complements NetAct-based GRN inference in elucidating the dynamical behaviors of the inferred GRNs. Together these two methods can be applied to infer biologically relevant regulatory interactions and the dynamical behavior of biological processes.

## Discussion

In this study, we have developed NetAct – a computational platform for constructing and modeling core transcription factor (TF)-based regulatory networks. NetAct takes a data-driven approach to establish gene regulatory network (GRN) models directly from transcriptomics data and takes a mathematical modeling approach to characterize cellular state transitions driven by the inferred GRN. The method specifically integrates both literature-based TF-target databases and transcriptomics data of multiple experimental conditions to accurately infer TF transcriptional activity based on the expression of their target genes. Using the inferred TF activity, NetAct further constructs a TF-based GRN, whose dynamics can then be evaluated and explored by mathematical modeling. Our approach in combining top-down and bottom-up systems biology approaches will contribute to a better understanding of gene regulatory mechanism of cellular decision making. NetAct is made freely available as an R package^71^.

One of the key components of NetAct is a pre-compiled TF-target gene set database. Here, we have evaluated different types of TF-target databases in identifying knocked down TFs using publicly available transcriptomics data sets. In this test, we have considered databases derived from literature, gene co-expression, cis-motif prediction, and TF-binding motif data. Our benchmark tests suggest that the literature-based database clearly outperformed the other databases. The literature-based database usually contains a small (∼30) number of target genes for each TF, but these data have direct experimental evidence, therefore being more reliable than those from the other sources. However, the literature-based database for sure has missing regulatory interactions, therefore maybe limiting the overall performance of NetAct. One way to address this issue is to further update the literature-based database, once new information is available. Another potential approach is to compile a database by combining different types of databases together. However, this might be quite challenging as different databases have data of very different sizes (the number of target genes) and quality. Future investigations on this direction can help to expand our knowledge of transcriptional regulation and meanwhile improve the performance of the algorithm.

NetAct also has a unique approach to infer the TF activity from the gene expression of the target genes with the consideration of activation/inhibition nature. From our in-silico benchmark tests, we found that NetAct outperforms major activity inference methods, owing to the design of the filtering step and the use of a high-quality TF-target database. NetAct is also robust against some inaccuracy in the TF-target database and noises in gene expression data, because of its capability of filtering out irrelevant targets as well as remaining key targets.

One potential issue is the assignment of the sign of TF activity, as it is algorithmically assigned according to the correlation with TF expression. In the case where the TF expression is very noisy or the expression is completely unrelated to TF activity, the sign assignment might be inaccurate. To deal with this issue, we have devised a semi-manual approach that identifies the sign of TF activity according to the sign of other interacting TFs. Another potential issue is that some TFs from the same family may have very similar target genes, therefore NetAct will have difficulty in identifying exactly which TF from the family is most relevant. Additional data resources, such as epigenomics^72^, TF-binding data^36^ and Hi-C data^73^, will be helpful to address this problem. One of the future directions is to design methods to integrate these data resources.

Lastly, instead of constructing a global transcriptional regulatory network, NetAct focuses on modeling a core regulatory network with only interactions between key TFs. The underlying hypothesis is that these TFs and the associated regulatory interactions play major roles in controlling the gene expression of different cellular states and the patterns of state transitions. With the core network identified using NetAct, we can further perform simulations with mathematical modeling algorithms, such as RACIPE, to analyze the control mechanism of the core network. These simulations allow us to generate new hypotheses, which can be further tested experimentally. The validation data can further help to improve the model. Ideally, this needs to be an iterative process to refine a core network model, which is indeed another interesting future direction.

## Conclusions

We developed NetAct, a computational platform for constructing and modeling core transcription-factor regulatory networks using both transcriptomics data and literature-based transcription factor-target gene databases. Utilizing both types of resources allows us to identify regulatory genes and links specific to the data and fully take advantage of the existing knowledgebase of transcriptional regulation. Our method in combining top-down and bottom-up systems biology approaches contributes to a better understanding of the mechanism of gene regulation driving cellular state transitions.

## Methods

### Selecting Enriched TFs

For a comparison between two experimental conditions, we obtained a ranked gene list quantified by the absolute value of the test statistics (t statistics in microarray and Wald test statistics in RNA-Seq) from differential expression (DE) analysis^74^, followed by gene set enrichment analysis (GSEA)^75^ using our optimized transcription factor (TF)-target gene set database. Here, for each TF, the corresponding gene set consists of all its target genes. GSEA identifies important TFs whose targets are enriched in DE genes between the two conditions. The significance test is achieved through 10,000 permutations of the gene list names and TFs are kept for further analysis when q value is below a certain threshold cutoff (0.05 by default). A C++ implementation of this version of GSEA, specifically for gene name permutations, has been provided in NetAct for fast computation. For multiple comparisons, a set of enriched TFs are first identified from each pairwise comparison and then a union of the multiple sets of TFs is considered.

In the database benchmark test, for each database, we computed the sensitivity and specificity values for different q-value cutoffs. Here, for each cutoff value, we defined the sensitivity as the proportion of data sets where the gene sets for the KD TFs were enriched with q-values below the cutoff value. We also defined specificity as the fraction of cases where the gene sets for the other TFs (non-KD TFs in the benchmark) were not enriched with q-values above the cutoff value. We then computed area under the ROC curve (AUC) using the DescTools R package^76^.

### Inferring TF activity

TF activity is inferred from the expression of target genes retrieved from the TF-target database. NetAct defines the activity of the selected TFs using two different schemes – one using only the expression of target genes and the other using the expression of both the TF and its target genes. The second scheme is only used for the situation of noisy target gene expression. For each TF, the algorithm selects the better scheme according to their performance, as described below.

#### Without directly using TF expression

For each TF, its downstream targets are first divided into two modules using the Newman’s community detection algorithm^77^ on the pairwise Spearman correlation matrix of the target genes. Then, within each module some less-correlated genes are filtered out to improve the quality of the inference. Here, the filtering step is achieved as follows: (1) each target gene is assigned a vector of correlations with the other target genes, where the distance between two genes is calculated as the sum of squares of the correlation vectors of two genes. (2) k-mean algorithm (k = 1) is performed within each cluster to determine the center vector. (3) genes are filtered out if the distance between the genes and the center is larger than the average distance.

This step outputs two groups of genes – genes in one group are supposed to be activated by the TF, while genes in the other group are inhibited by the TF. Note, at this stage, the nature of activation/inhibition of the individual group is not yet determined. The activity of the TF is calculated as

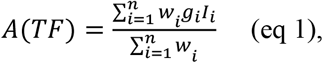

where *g*_*i*_ is the standardized expression value of a target gene *i, w*_*i*_ is the weighting factor defined as a Hill function:

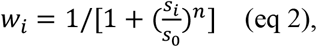

where *s*_*i*_ is the adjusted p value from DE analysis for gene *i*, the threshold *S*_0_ is 0.05, and n is set to be 1/5 for best performance (Fig. S8). *I*_*i*_ is 1 if the corresponding gene belongs to the first group and -1 if it belongs to the second group. If the calculated TF activity pattern is not consistent with the TF expression trend (evaluated by Spearman correlation), both the sign of the two groups and the sign of the activity are flipped.

According to our in-silico benchmark test (Fig. S9), we found that majority of the targets in one group are activated by the TF, and majority of those in the other group are inhibited by the TF. For genes in the inhibition group, the higher the TF activity, the more the genes are suppressed. Thus, the formula in Equation (1) captures well the activity of TFs for their effects to both activating and inhibitory targets. We also explored a few other community detection algorithms^78–80^ and found they produced similar results (Fig. S1).

#### Using TF expression

For each TF, its downstream targets are first divided into two groups according to the sign of the Spearman correlation between the TF expression and the target expression. Similar to the previous scheme, in each group, target genes are filtered out if the correlation value is less than the average correlation of all the targets. The activity of the TF is also calculated using Equation 1.

#### Sign assignment for DE TF

For any DE TF (*i*.*e*., there is significant difference in TF expression across cell type conditions) of interest, NetAct computes the activity values from both the schemes (with or without TF’s expression), and selects the better way based on how well the activity values correlate with target expression. To this end, NetAct calculates the absolute value of Spearman correlation between the TF activity and the expression of each target, and selects the scheme whose activity gives larger average correlations.

#### Sign assignment for non-DE TF

If the expression patterns of the identified TFs fail to show the significant differences between cell type conditions, a semi-manual method to assign the sign of activity can be adopted. Putative interaction partners between DE and non-DE TFs in the inferred network are identified using the Fisher’s Exact Test between TF targets in the NetAct TF-target database. The most significant pairs are then cross referenced with the STRING database to identify instances of PPI. A literature search is then performed to identify the nature of the PPI, and the sign of the non-DE TF is adjusted based on the DE TF and the type of PPI. Note that the last step needs to be done manually for each modeling application. Table S3 shows the details of TF sign flipping and supported experimental evidence for the two network modeling applications.

### Network construction and mathematical modeling

NetAct constructs a TF regulatory network using both the TF-TF regulatory interactions from the TF-target database and the activity values. (1) The network is constructed using mutual information between the activity values of two TFs. (2) Interactions are filtered out if they cannot be found in the TF-target regulatory database (*i*.*e*., D1). (3) The sign of each link is determined by the sign of the Spearman correlation between the activity of two TFs. (4) We keep the interaction between two TFs if their mutual information is higher than a threshold cutoff. With different cutoff values for mutual information, NetAct establishes networks of different sizes. To identify the best network model capturing gene expression profiles, we apply mathematical modeling to each of the TF networks using RACIPE^29^. RACIPE takes network topology as the input and generates an ensemble of mathematical models with random kinetic parameters. By simulating the network, we expect to obtain multiple clusters of gene expression patterns that are constrained by the complex interactions in the network. RACIPE was also applied to generate simulated benchmark test sets for a synthetic TF-target network (see Supplemental Material SI5).

## Declarations

### Ethics approval and consent to participate

Not applicable

### Consent for publication

Not applicable

## Availability of data and materials

The information of the TF-target gene set databases is listed in Table S1. The public gene expression datasets for algorithm optimization and benchmark are listed in Table S2. The datasets and computational scripts for in-silico benchmark, the network modeling scripts, including those for data processing, network construction and network simulations, and the inferred network topology files are available in the GitHub repository at https://github.com/lusystemsbio/NetActAnalysis. The NetAct software is available at https://github.com/lusystemsbio/NetAct as an R package. NetAct is platform independent, written in R with a partial of codes in C++ for improved performance. NetAct is licensed under the MIT License.

## Competing interests

The authors declare that they have no competing interests

## Funding

The study is supported by startup funds from The Jackson Laboratory and Northeastern University, by the National Cancer Institute of the National Institutes of Health under Award Number P30CA034196, and by the National Institute of General Medical Sciences of the National Institutes of Health under Award Number R35GM128717.

## Authors’ contributions

M.L conceived the study. K.S. developed and V.K. and A.K. improved the NetAct algorithm. A.K. constructed and performed in-silico benchmark. K.S. and V.K. performed benchmark tests on public experimental gene expression data. B.C. and V.K. performed network modeling. D.D. helped to refine the NetAct code. S.L, K.K., and Z.S.Q. provided conceptual input to the manuscript. K.S., A.K., V.K. and M.L wrote the manuscript, with helps from all other authors. The authors read and approved the final manuscript.

## Acknowledgements

Not applicable

## Notes

### Competing Interest Statement

The authors have declared no competing interest.

### Summary of Updates

main text and figures, SI text, SI figures

