## Supplemental Text for "NetAct: a computational platform to construct core transcription factor regulatory networks using gene activity"

### Supplemental Information

#### 1. TF-target gene set databases

We constructed transcription factor (TF)-target databases for both human and mouse genomes by using several existing databases of different types, as listed in Table S1. Effectively, we constructed databases of four types, namely, D1, D2, D3, and D4. The D1 type consists of literature-based TF-target relationships, including (a) TTRUST [1], a TF-target interaction database using text mining; (b) RegNetwork [2], a data repository of gene regulations, where we incorporated data from both experiments and predictions with high confidence; (c) TFactS [3], a predictive regulatory database based on microarray experiments; (d) TRED [4], a precise trans-regulatory database for target genes of cancer-related TFs. For the D2 type, we merged 394 tissue-specific gene regulatory networks from FANTOM5[5] to form a weighted target database for 643 TFs. The FANTOM5 networks were inferred from gene expression data of various tissues. The D3 type includes public databases of experimentally-validated eukaryotic TFs and their target genes, TRANSFAC[6] and JASPAR[7], a database derived from ChIP-X experiment, ChEA[8], and a database that tracks sthe binding regions of the TFs from various ChIP-seq experiments, ENCODE[9]. For the D4 type, we considered RcisTarget[10], which identifies TF-target gene relationships based on TF binding motifs. RcisTarget includes both literature and prediction data.

All the gene names from these databases are corrected using genome wide annotation databases (org.Hs.eg.db and org.Mm.eg.db, for additional org.xx.eg.db packages, see *http://www.bioconductor.org/packages/release/data/annotation/*), GeneCards for human genes, and Mouse Genome Informatics (*http://www.informatics.jax.org/*) for mouse genes. For future expansion, we provide two versions of the databases – one with the format of gene symbols and the other with the format of Entrez Gene ID[11]. Detailed description of all these databases, such as the total number of covered TFs, average number of targets for each TF, is listed in the Table S1.

#### 2. Processing transcriptomics data

NetAct takes gene expression profiles as input. Each gene profile corresponds to a grouping scheme and a design of comparisons. For microarray data, the transcriptomics data were pre-processed using a standard protocol -- the raw data files (.CEL) were processed by affy[12] with robust multichip average (RMA) normalization[13]. For RNA-Seq data, we took raw count matrix and converted them into the measurement of log counts per million (log-cpm).

Typical transcriptomics profiles contain data of two or more experimental conditions, *e.g.*, a control group (C) with replicates ($C_1$, $C_2$, *etc.*) and a treatment group ($T_1$, $T_2$, *etc.*). To prepare for further downstream analysis, NetAct takes user-input of the grouping scheme and the design of comparisons and performs all requested differential expression (DE) analyses. This step is conducted by limma package[14] on microarray data and DESeq2[15] (we also provide an option of limma + Voom[16], *e.g.*, as we used in network modeling) on RNA-Seq raw count. For a pairwise comparison, *e.g.*, group C versus group T, both limma and DESeq2 output test statistics and adjusted p-values; while for cases with multiple conditions: with limma, we first perform individual pairwise comparison and then obtain F statistics and associated adjusted p-values; with DESeq2, we perform the likelihood ratio test to acquire the corresponding statistics and associated adjusted p-values.

#### 3. *In silico* TF activity inference benchmark

We benchmarked NetAct activity inference against other three activity inference methods, namely, FastNCA[17], Viper[18], and AUCell[10], each method with distinct approaches to infer the TF activity. (i) FastNCA: The MATLAB code is available at https://www.eee.hku.hk/~cqchang/FastNCA.htm. The program takes two inputs the expression matrix and the connectivity matrix constructed from the regulons to the TF activity. (ii) Viper: The R package is available at https://www.bioconductor.org/packages/release/bioc/html/viper.html. The viper function in the R package uses the target gene expressions and the regulons to calculate the TF activity. (iii) AUCell: The R package is available at https://bioconductor.org/packages/release/bioc/html/AUCell.html. The getAUC function in the R package uses the cell rankings obtained from the expression matrix and the regulons to calculate the TF activity. We obtained the expressions matrix by simulating a synthetic gene regulatory network (GRN) which also gives us the regulons, *i.e.*, the TF-target gene relationships.

#### 4. Construction of the synthetic GRN

We took the following approach to create the synthetic gene regulatory network (GRN) to mimic a TF-target gene network where the interactions between a set of TFs and their target genes drive the downstream processes. First, we created a set of 30 TFs, labeled as tf1, tf2, tf3, …, tf30. We established interactions among these 30 TFs by randomly assigning two links for each TF, permitting self-interactions. Each TF to TF interaction type could be one of excitatory, inhibitory, or signaling, with a probability of 25%, 25%, and 50%, respectively. In one of the realizations of this assignment, 25 out of these 30 TFs are the targets of the other TFs or themselves. Second, we created a pool of 1,000 target genes, labeled as tg1, tg2, tg3, …, tg1000. Then, for each TF, we randomly assigned 20 genes from the target pool as the TF targets. The TF to target interaction type could be either excitatory or inhibitory with a 50% chance for each type. The TF-target network is illustrated in Fig. 3a. The TFs are shown in large yellow filled circles and the targets are in small blue filled circles. In total, the simulated network has 477 nodes including 30 TFs and 447 target genes. The signaling interactions affect the activities, not the expressions. The constructed network has a total of 660 interactions and 13,650 non-interacting pairs. Also, the labeled transcription factor TF9 in the figure was selected for knockdown simulations.

**Construction of unperturbed and perturbed regulon database**

We built a regulon database (DB) which is the collection of all TF to target gene interactions. While listing the target genes of a TF, we consider the excitatory and inhibitory interaction types only, *i.e.*, we excluded the target genes with the signaling interaction. As a result, for a certain TF, the number of targets could be 20 (when both TF to TF interactions are signaling), 21 (when one of the TF to TF interactions is signaling), or 22 (when none of the TF to TF interactions are signaling).

An important question is how a noisy TF-target database can influence the accuracy in predicting TF activity. To be able to explore this question, we generated perturbed TF-target databases for a more challenging benchmark. From the regulon database created above, we derived 100 perturbed regulon databases at each of the following perturbation levels: 25%, 50%, and 75%. To construct a perturbed regulon DB, we initially created a target pool by including all the targets that are not TFs in the unperturbed regulon DB. Subsequently, for each TF, we replaced a subset of the targets that are not TFs by other targets from the target pool. The number of the replaced targets is determined by the perturbation level. To perform this replacement, we first randomly selected and removed required number of targets from the target set of a TF. Then, we randomly sampled (without replacement) the same number of targets from the target pool that is created above. We placed the selected targets into the target set of the TF. We repeated the above process for each TF to construct a perturbed regulon DB at a specific perturbation level. Furthermore, we created a second version of the above unperturbed and perturbed regulons to satisfy the AUCell protocol, which advises to include the target genes with only positive interactions in the regulons. To modify the unperturbed regulons, we retained the targets with the excitatory TF-target regulation. On the other hand, to modify the perturbed regulons, for each TF, targets fall into two groups: unperturbed targets (the targets that were also in the unperturbed regulon), and perturbed targets (the targets that got incorporated because of random perturbation). New targets for a TF will incorporate the perturbed targets with positive TF-target relationship and a random half of the perturbed targets, assuming that half of the target genes are positively regulated (denoted as AUCell 1). AUCell was also applied using the same regulons (both unperturbed and perturbed regulons) as those used in the other three methods (denoted as AUCell 2).

#### 5. Simulation of activity and expression using RACIPE

We utilized our previously develop mathematical modeling method, Random Circuit Perturbation (RACIPE)[19], to simulate the expression of network genes and the activities of the TFs. We first consider a network with only transcriptional regulation. In RACIPE, for a gene T that is transcriptionally regulated by multiple regulators $R_i$ $(i = 1, 2 \ldots)$, the gene expression dynamics is given by an ordinary differential equation (ODE)

$$de_T/dt = \frac{G_T}{\prod_i \lambda^{+}{}_{R_iT}} \prod_i H^S(e_{R_i}, R_iT_0, n_{R_iT}, \lambda_{R_iT}) - k_T e_T \quad , \qquad \text{(S1)}$$

where $e_T$ and $e_{R_i}$ are the gene expression levels of genes T and $R_i$, respectively, $G_T$ is its maximum production rate, and $k_T$ is the degradation rate of gene T. $H^S$ is the shifted hill function used for a regulatory interaction from R to T and is given by,

$$H^S(e_R, RT_0, n_{RT}, \lambda_{RT}) = \lambda_{RT} + (1 - \lambda_{RT})/(1 + \left(\frac{e_R}{RT_0}\right)^{n_{RT}}) \ . \qquad \text{(S2)}$$

Here $RT_0$, $n_{RT}$ and $\lambda_{RT}$ are the threshold level, the Hill coefficient of regulation, and the maximum fold change for the regulatory link from R to T. For an excitatory interaction, $\lambda_{RT}$ is denoted as $\lambda^{+}{}_{RT}$ and is greater than one, and $H^S$ ranges from (1, $\lambda^{+}{}_{RT}$); while for an inhibitory interaction, $\lambda_{RT}$ is denoted as $\lambda^{-}{}_{RT}$ and is smaller than one, and $H^S$ ranges from $(\lambda^{-}{}_{RT}, 1)$. The term $\prod_i \lambda^{+}{}_{R_iT}$ in Eq. S1 denotes the product over all excitatory interactions of T and serves as a scaling factor to ensure $G_T$ as the maximum production rate. Using this formalism, RACIPE generates models with kinetic parameters randomly selected from uniform distributions, *i.e.*, $G_T$ from (1, 100), $k_T$ from (0.1, 1), $n_{RT}$ (integer) from (1, 6), and $\lambda^{+}{}_{RT}$ from (1, 100). For $\lambda^{-}{}_{RT}$, RACIPE samples from a uniform distribution of (1, 100) and then takes the inverse. $RT_0$ is taken from ($0.02M$, 1.98M), where the median Hill threshold M is estimated according to the half-functional rule[20]. For each randomly generated model, RACIPE simulates the ODE for the whole network (Eq. S1 as an example for gene T) starting from an initial condition of $e_T$ randomly selected from a logarithmic distribution whose maximum is $\frac{G_T}{k_T}$, and minimum is $\frac{G_T}{k_T}(\frac{\prod_i \lambda^{-}{}_{R_iT}}{\prod_i \lambda^{+}{}_{R_iT}})$. From an ODE simulation of each model, we evaluate whether the steady-state solution exists and, if so, obtain the steady-state gene expression profile.

For a network with both transcriptional and signaling regulations, we adopted a generalized RACIPE approach, as described in a previous study[19]. For a gene T (technically T needs to be also a regulator to be considered in regulator's activity calculation) that is transcriptionally regulated by multiple regulators $R_i$ ($i = 1, 2 ...$) and is regulated by signaling regulation by $S_j$ ($j = 1, 2 ...$), the gene activity dynamics of T is given by an ODE

$$da_T/dt = \frac{G_T}{\prod_i \lambda^+{}_{R_iT} \prod_j \lambda^+{}_{S_jT}} \prod_i H^S(a_{R_i}, R_iT_0, n_{R_iT}, \lambda_{R_iT}) \prod_j H^S(a_{S_j}, S_jT_0, n_{S_jT}, \lambda_{S_jT}) - k_T a_T, \quad \text{(S3)}$$

where $a_T$, $a_{R_i}$ and $a_{S_j}$ are the gene activity levels of genes T, $R_i$, and $S_j$, respectively. Here, in the generalized RACIPE, regulator's activity is modelled phenomenologically (instead of Michaelis-Mention kinetics) by using Hill functions to capture ultra-sensitivity commonly observed in signaling regulation[21]. The generalized RACIPE has been applied to model a cell cycle gene regulatory network[19]. Meanwhile, we model the gene expression dynamics of T by

$$de_T/dt = \frac{G_T}{\prod_i \lambda^+{}_{R_iT} \prod_j \lambda^+{}_{S_jT}} \prod_i H^S(a_{R_i}, R_iT_0, n_{R_iT}, \lambda_{R_iT}) - k_T e_T. \quad \text{(S4)}$$

Here, the signaling regulation by $S_j$ affects gene activity but not gene expression.

RACIPE is able to generate two set of the activities for two cases: a *wide-type* scenario with untreated samples, and a *knockdown* scenario with both untreated samples and samples with gene knockdown (specifically TF9 in the test). For the first scenario, we performed RACIPE simulation to calculate both the expressions and the activities of the genes in the synthetic GRN. From randomly generated RACIPE models, we found 83 models that generate stable steady states. For the second scenario, we performed knockdown simulations in which selected genes to be expressed minimally. Here, we lowered TF9's production rate by 95%. Similar to the wild type case, we performed RACIPE simulations to calculate both the activities and expressions of the network genes[22]. From 100 randomly generated RACIPE models, we found 86 models that generate stable steady states. Furthermore, we combined both the controls and treatments of a subset of 40 models from the wild type simulations with those of a subset of 43 models from the knockdown simulations.

**6. *In silico* network construction benchmark**

We compared NetAct with three other existing top-performing methods GENIE3[23], GRNBoost2[24], and ppcor[25,26] for network inference. In this benchmark test, we used the same *in-silico* data generated for the TF activity benchmark. The benchmark data set contains the ground-truth TF-target interactions obtained from the synthetic GRN (see SI4) and the gene expression and TF activity profiles of 83 models for the GRN simulated with RACIPE (see SI5). We used the expression profiles as the input for network construction and evaluated the performance against all the ground-truth TF-target interactions.

For the NetAct implement, as the method requires a TF-target regulatory database (regulon database), we generated the database corresponding to the original regulons from the synthetic GRN. In this test, as the TF-target relationships were provided to NetAct as an input, the purpose of the test is to evaluate how well the NetAct-inferred TF activity can be used for network construction to recapitulate the same TF-target interactions. The details of the NetAct testing procedure are described as follows. First, we inferred regulatory interactions from TFs to target genes by applying NetAct using the simulated gene expression data and the original regulon database. Note that NetAct first infers TF activity before network construction. Second, we compared the inferred interactions with the ground-truth interactions to calculate the precision and recall at different mutual information (MI) thresholds, which were obtained from the distribution of the MI values. When performing this comparison, we consider an interaction to be correct when both the regulator and target genes are presented in a ground-truth interaction, regardless of which gene is the regulator. NetAct can learn the directions of interactions from regulons, but we chose not to evaluate them in the benchmark, as the directionality is not considered in other network construction algorithms.

We also evaluated the performance of NetAct in inferring regulatory interactions with regulons at different perturbation levels (25%, 50%, and 75%, same as the conditions in the TF activity benchmark). The purpose of this test is to evaluate how the performance of NetAct is affected when the regulon database is noisy or inaccurate. In this test, we first applied NetAct to infer TF activities using a perturbed regulon database, followed by network construction using mutual information. For each perturbation level, we evaluated the precision and recall values by (1) comparing the inferred interactions with all the ground-truth interactions in the synthetic GRN and (2) comparing the inferred interactions with the ground-truth interactions, excluding those that are presented in the perturbed regulon database. In the latter case, we do not consider the interactions in the perturbed regulon database to emphasize on the prediction of novel interactions.

We also evaluated the performance of the other three network construction algorithms GENIE3, GRNBoost2, and ppcor using the simulated gene expression profiles. GENIE3 and GRNBoost2 takes gene expressions and the list of TFs as the input; then they predict interactions from the designated TFs to the target genes together with an importance score for each interaction. We calculated the precision and recall at different importance score thresholds, which were obtained from the distribution of the importance scores. Ppcor takes the gene expressions as the input and predicts gene to gene relationship together with signed correlation values. The method does not treat TFs separately from their targets. We retained the interactions whose sources are TFs and calculated the precision and recall for the network inference at different absolute correlation thresholds, which were obtained from the distribution of the absolute correlation values. Additionally, to mimic SCENIC[10], we applied GENIE3 to the activity profiles calculated using AUCell from the simulated gene expression profiles and calculated the precision and recall as before (GENIE3+AUCell).

**7. Applications of network modeling with NetAct**

For modeling the EMT network, we downloaded the microarray gene expression data from GSE17708. Gene symbols were mapped using R package hgu133plus2.db[27]. Unannotated genes were removed and mean expression was used for redundant probes. Samples were split into three groups – "early" for data from untreated, 0.5h and 1h, "middle" for data from 2h, 4h and 8h, and "late" for data from 16h, 24h and 72h. A three-way comparison (early-middle, middle-late, early-late) was performed for differential expression (DE) analysis using limma[28] (in NetAct, we provide the function MultiMicroDegs for a multi-way comparison). Enriched transcription factors (TFs) were identified using a modified algorithm of gene set enrichment analysis (GSEA)[29] (in NetAct with the function TF_GSEA) for each of the three comparisons. Significantly enriched TFs were identified from the GSEA with a q value cutoff of 0.01, and the identified TFs from the three comparisons were combined. Next, activity of enriched TFs was inferred using the function TF_Activity in NetAct, followed by the construction of a = TF regulatory network using TF_Filter (miTh = 0.05, nbins = 8, cormethod = "spearman", DPI = T). To assign the sign of activity for non-DE TFs, we performed the fisher exact test (see Methods) and cross referenced with stringDB[30] to identify the putative interaction partners between the non-DE and DE TFs (Table S3A, 1$^{st}$ and 2$^{nd}$ column). We then performed literature search to check whether the two TFs interact synergistically or an antagonistically (Table S3A, 3$^{rd}$ – 4$^{th}$ column), and checked whether the signs of the two TFs agree with the nature of the interactions or not. If not, the sign of activity for the non-TF was flipped (Table S3A, 5$^{th}$ column). The final topology generated was then simulated with sRACIPE[31] for deterministic analysis using 10,000 models and default parameters. TF KD analysis was performed using the function sracipeKnockDown function in the sRACIPE R package with nClusters = 2.

For modeling the TF regulatory network for macrophage polarization, the counts matrix was obtained from GSE84517, and duplicated gene IDs were removed. Gene IDs were converted to symbols using the function select ( org.Mm.eg.db, columns="SYMBOL", keytype="ENSEMBL") from the R package AnnotationDbi[32]. Count data was then filtered to remove those with less than 10 reads in all samples. DE results were generated using Limma and Voom (?) (MultiRNAseqDegs_limma from the NetAct R package) comparing UT-IL4_2h, UT-IFNg_2h, UT-IFNg_IL4_2h, UT-IL4_4h, UT-IFNg_4h, UT-IFNg_IL4_4h. Batch effects were adjusted using ComBat[33]. Enriched TFs were identified for each comparison using the NetAct function TF_GSEA (minSize=5, nperm = 10000, qval = T). For all comparisons except UT-IL4_2h, a q-value cutoff of 0.01 was used to identify enriched transcription factors which were then combined. For UT-IL4_2h, a relaxed q-value cutoff of 0.05 was used in order to increase the number of TFs that show differential activity in the UT vs IL4 states. Without the relaxed q-value cutoff only very few TFs showed unique expression patterns between the UT and IL4 states. Enriched TFs were then combined, and activity was inferred using the NetAct function TF_activity (with_weight = TRUE, useDatabaseSign = F, useCorSign = T, if_module = F). Three networks were then constructed all using the NetAct function TF_Filter (miTH = 0.4, nbins =3, maxTf = 150, maxInteractions = 500, corMethod = "s", useCor = T, removeSignalling = FALSE, DPI = F, miDiff = 0.0). These networks were combined and discrepancies in edges were decided by the most common edge, with ties set to activating. The assignment of the sign of activity for non-DE TFs is described in Table S3B.

The final TF network topology was then simulated using sRACIPE for deterministic analysis with 10,000 models and default parameters. Each resulting model was mapped back to experimental states by first averaging the experimental values of the simulated genes in each experimental condition and then identifying the experimental condition with the lowest Euclidean distance to each model. The color schemes of the mapping can be observed in Fig. 6f for the simulated heatmap and in Fig. 6e for the data projected to the first two PCs. The structure of the projected simulation data is very close to the structure of the projected experimental data. Moreover, we observed consistent patterns of gene expression/activity in the experimental and simulation data. For example, we found high Sp1/3 and Foxo3 in the UT condition but low in other conditions, high Stat1 and Nfkb1 in the IFNγ and IFNγ +IL4 conditions, high Rb1 and Smad1 in the UT and IL4 conditions, and high Myc and Ep300 in the IL4 condition. These observations are largely consistent with the literature[34].

To analyze the transition paths, we started by projecting simulation gene expression data of the macrophage network to the first two principal components and separating the data with two lines ($PC2 = 2PC1 - 11$ and $PC2 = 2PC1$ (blue, green, and red in Fig. S10A). The group in green in Fig. S10A corresponds to the transitory models (Fig. S10B for models corresponding to different treatment conditions). We then computed the orthogonal projection of the expression data of the transitory models to the middle of the two lines ($PC2 = 2PC1 - 5.5$) (Fig. S10C). The histogram of the points on the regression line were analyzed to determine transition path behavior (Fig. S10D).
