## Supplemental Figures for "NetAct: a computational platform to construct core transcription factor regulatory networks using gene activity"

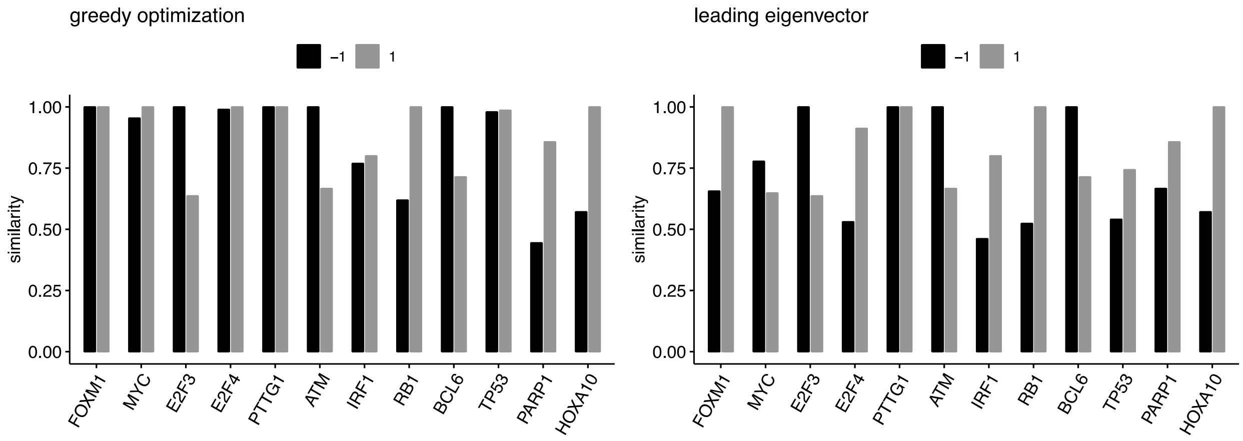


**Fig. S1: Comparison of the grouping schemes by Newman's method (NetAct) and other community detection algorithms.** Two other community detection algorithms were applied to divide target genes of enriched transcription factors into two groups using the gene expression data from shRNA knockdown of FOXM1 in lymphoma cells (GEO: GSE17172). Here, we applied two other community detection algorithms: greedy optimization (left) and leading eigenvector (right) algorithms. Within the detected communities, we calculated the maximum percentage of genes overlapped with the gene members detected by NetAct (defined as similarity). As shown, the gene members from one community corresponding to the +/- regulatory direction can be largely overlapped with the grouping schemes from other community detection algorithms.

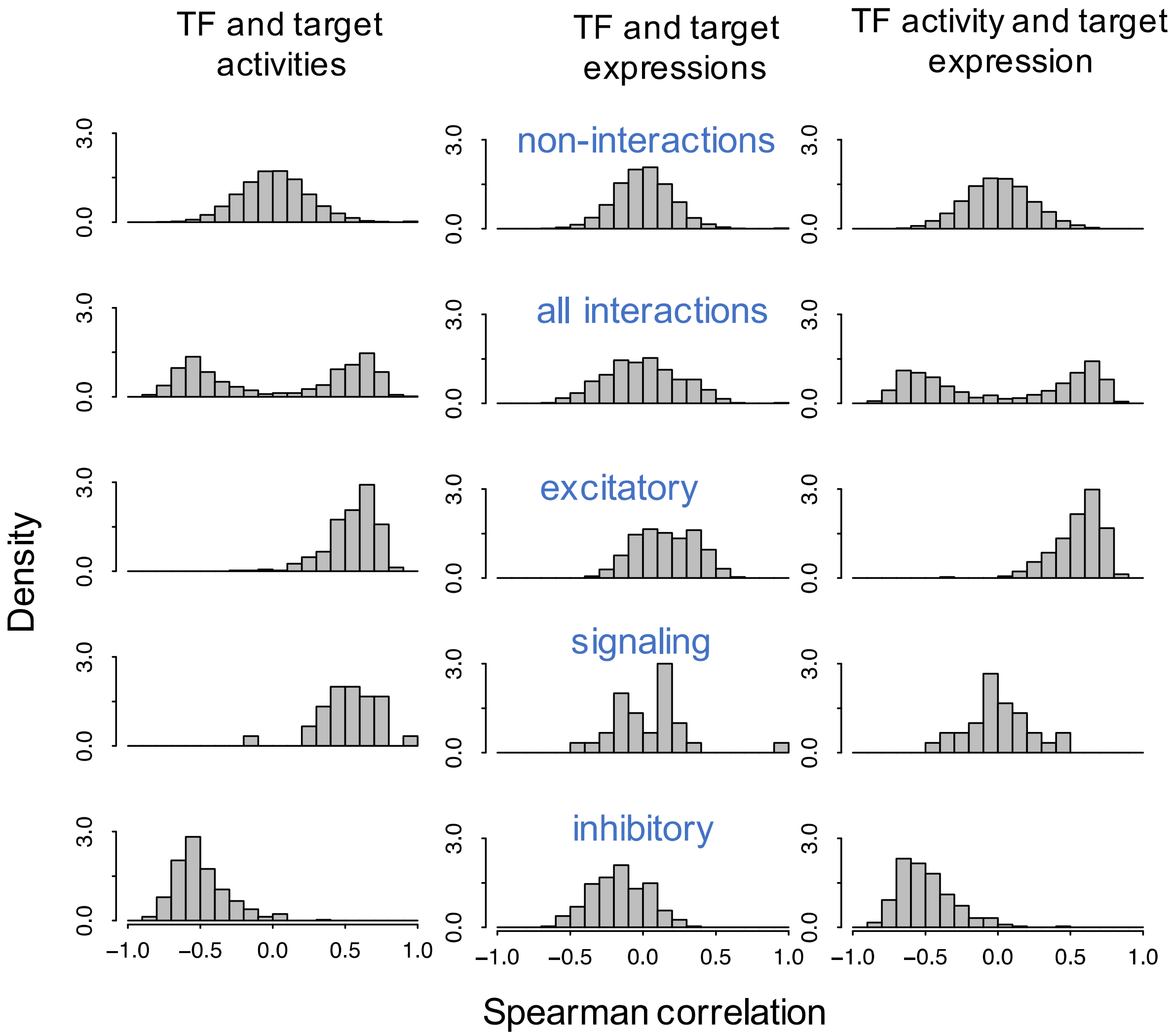


**Fig. S2. Correlation structure in the simulated activities and expressions of the synthetic gene regulatory network with knockdown of transcription factor TF9.** RACIPE generated simulated expressions of the synthetic gene regulatory network (shown on Fig. 3a) with knockdown of transcription factor TF9. Histogram of Spearman correlations between transcription factor (TF) and target gene activities (column 1), TF and target gene expressions (column 2), and TF activities and target gene expressions (column 3). In each case, different interaction types are shown in separate rows - row 1: non-interactions, row 2: all interactions, row 3: excitatory, row 4: signaling, and row 5: inhibitory.

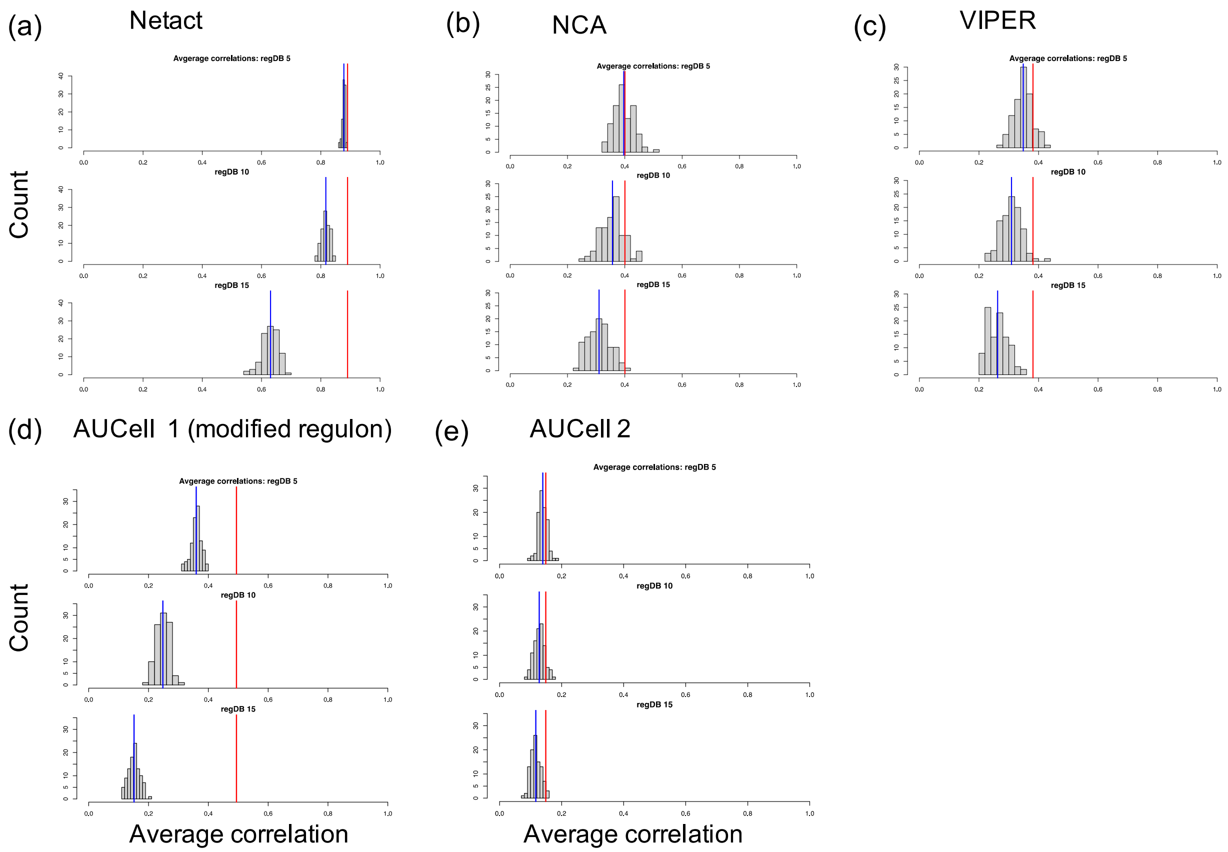


**Fig. S3. Comparing stability of activity inference methods.** (**a**) NetAct, (**b**) NCA, (**c**) VIPER, (**d, e**) AUCell. These four different methods applied to infer TF activities using RACIPE simulated expressions of the synthetic GRN (shown on **Fig. 3a**) and the regulons (obtained from the synthetic GRN). The regulons used at three levels of target gene perturbations: 25% (regDB 5), 50% (reg DB 10), and 75% (regDB 15). At each perturbation level, 100 instances of the perturbed regulons are created. For each instance of the perturbed regulons, NetAct and the three other methods are applied to infer TF activities from the RACIPE simulated TF expressions and Spearman correlations between the RACIPE simulated TF activities (ground truth) and inferred TF activities are calculated and then averaged. Histograms of the average Spearman correlations obtained using all 100 instances of the perturbed regulons at each perturbation level are shown here. The blue vertical line depicts the mean of the average Spearman correlations at each perturbation level. The red vertical line on each panel marks the average Spearman correlation using the unperturbed regulon (perturbation level 0%). AUCell requires that regulons include only the positive interactions and so we modified the regulons (both unperturbed and 100 perturbed instances at each perturbation level) to comply with AUCell's protocol. For TF activity calculation by applying the AUCell method, we used both regulons: these modified regulons and named the calculations as AUCell 1 (**d**) and the same regulons used in the three other methods (labelled as AUCell 2) (**e**).

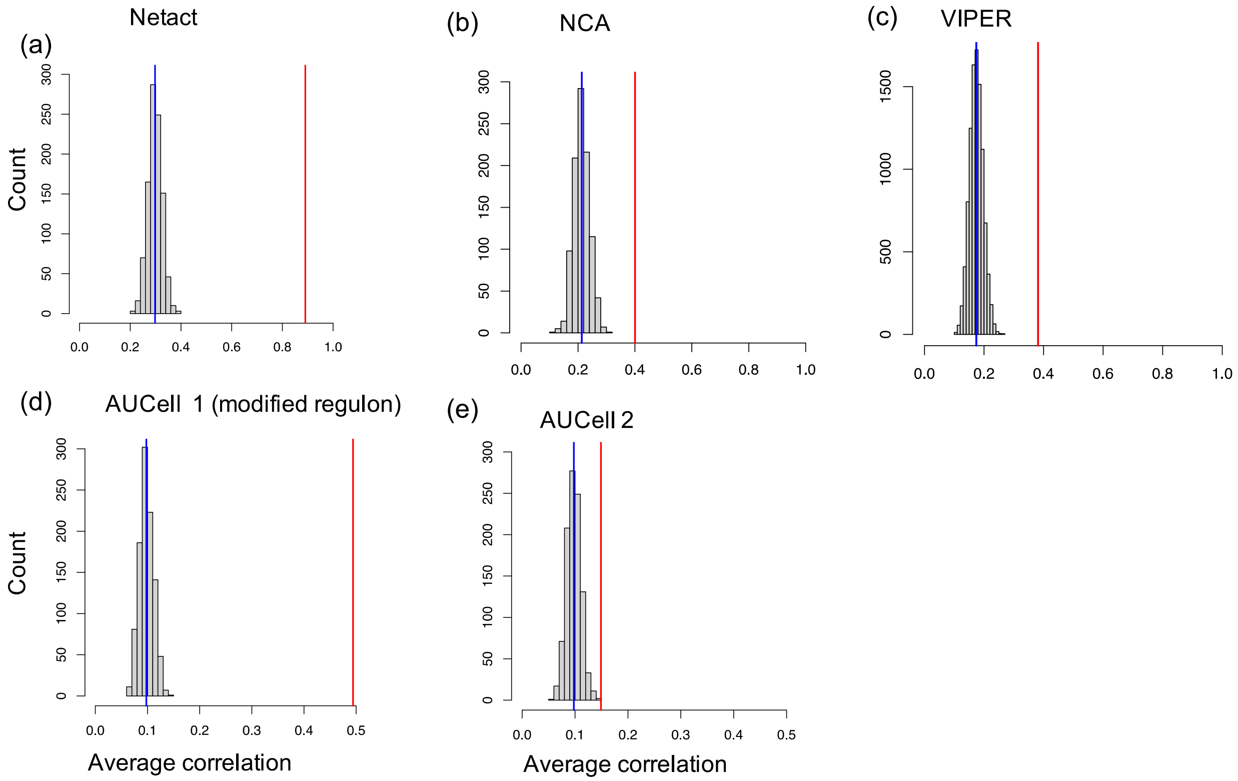


**Fig. S4. Null distribution of the average correlations for the four methods**. (**a**) NetAct, (**b**) NCA, (**c**) VIPER, (**d**) AUCell 1 (modified regulons), and (**e**) AUCell 2. First, RACIPE expressions and activities (ground truth) are calculated for the synthetic GRN (shown in **Fig. 3a**). Then, the gene labels of the RACIPE expressions are shuffled and the TF activity inference methods are applied to infer the TF activities using the unperturbed regulons. To calculate the average correlation, first Spearman correlations between the RACIPE activities (ground truth) and the inferred TF activities by a method are calculated and then averaged over across the TFs. The process is repeated 100 times to obtain the distribution of the average correlations. The red vertical line depicts the average correlation obtained from the RACIPE activities (ground truth, without gene shuffling) and the inferred TF activities. AUCell protocol requires that AUCell requires that regulons include only the positive interactions and so we modified the regulons to comply with AUCell's protocol. For TF activity calculation by applying the AUCell method, we used both regulons: these modified regulons and named the calculations as AUCell 1 (**d**) and the same regulons used in the other three methods (labelled as AUCell 2) (**e**).

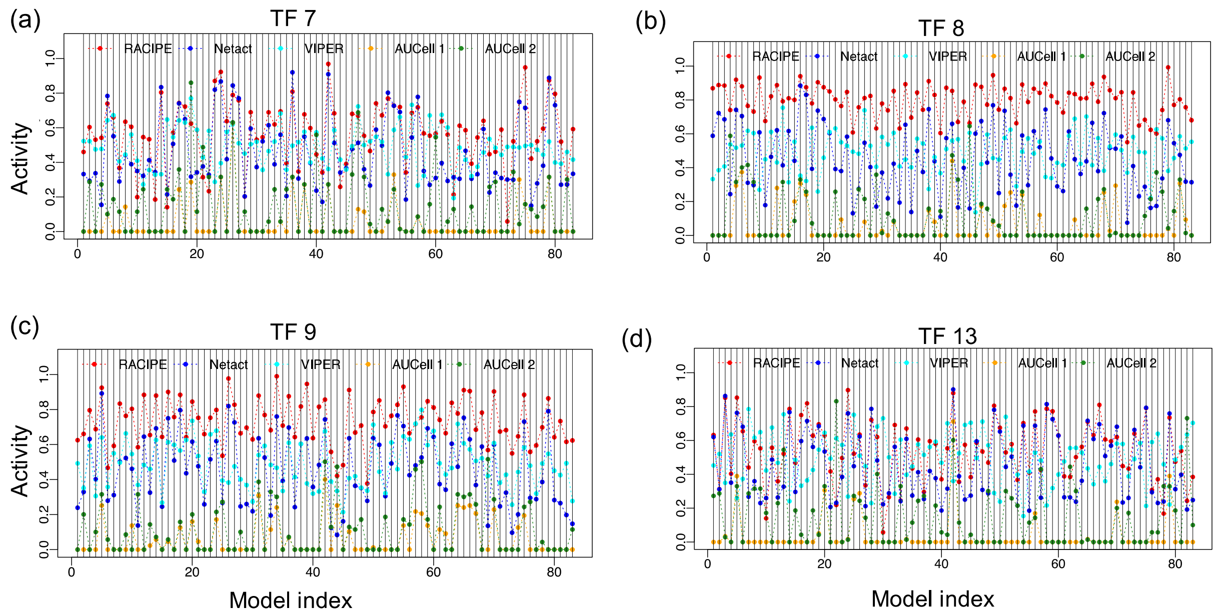


**Fig. S5. Activity levels of four transcription factors**. Normalized activities of four representative transcription factors (TFs) on the synthetic GRN (shown in **Fig. 3a**). (**a**) TF 7, (**b**) TF 8, (**c**) TF 9, and (**d**) TF 13. TF activities are inferred from RACIPE simulated expressions by using four methods NetAct, NCA, VIPER, AUCell 1 (using regulons modified according to AUCell protocol, see **Fig. S3** caption), and AUCell 2 (using the same regulons as used in NetAct, NCA, and VIPER).

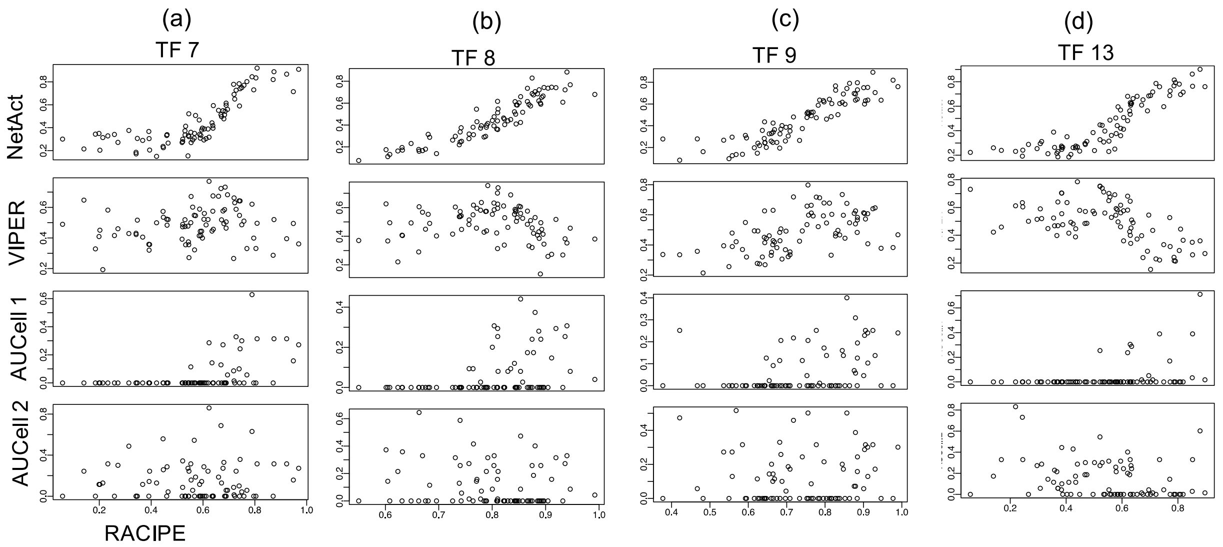


**Fig. S6. Scatter plot for activities of four transcription factors**. Scatter plot of normalized activities from RACIPE and each of the four inference methods NetAct, NCA, VIPER, AUCell 1 (using regulons modified according to AUCell protocol), and AUCell 2 (using the same regulons as used in NetAct, NCA, and VIPER). Four representative transcription factors (TF) are shown: (**a**) TF 7, (**b**) TF 8, (**c**) TF 9, (**d**) TF 13 on the synthetic GRN shown on **Fig. 3a**.

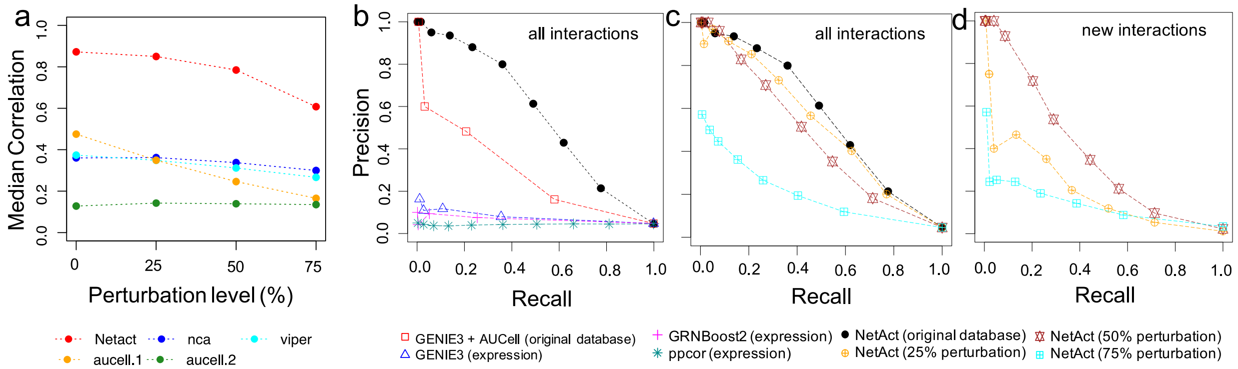


**Fig. S7. The performance of activity inference and network construction from a simulation benchmark**. The analyses are similar to those presented in Fig.4. But in this case, the benchmark was performed on the simulated data of the synthetic gene regulatory network (GRN) under the condition of TF9 knockdown. **(a)** TF activity inference. TF activity was inferred by several methods using the gene expression data simulated from the synthetic TF-target GRN and the corresponding regulons. For each TF, we computed Spearman correlations between the inferred activity and simulated activity (ground truth) for all the simulated models. Then, we calculated the average correlation values over all TFs. The plots show the median of average correlations for the cases where we used the original regulons defined by the TF-target network (0% perturbation), and the regulons where 5 (25% perturbation), 10 (50% perturbation), and 15 (75% perturbation) target genes are randomly replaced with non-interacting genes, respectively. The median values were computed over 100 repeats of random replacement for each perturbation level, and the values of the average correlations are reported for the case of zero perturbation. Shown are the results for NetAct (red), NCA (blue), VIPER (cyan), AUCELL 1 where regulons contain only positively associated target genes (orange), and AUCELL 2 where regulons contain all target genes (green). **(b-d)** Network inference. The panels show the performance of network inference algorithms from the simulation benchmark by the precision and recall for different link selection threshold. **(b)** Network inference performance against all ground-truth regulatory interactions. Tested methods are GENIE3, GRNBoost2, and PPCOR, using transcription factor (TF) expression; GENIE3 using TF activity inferred by AUCell; NetAct using its inferred TF activity. For the latter two methods, original (unperturbed) regulons were used. **(c)** Network inference performance of NetAct against all ground-truth regulatory interactions using the regulons with 0% (original regulons), 25%, 50%, and 75% target perturbations. **(d)** Network inference performance of NetAct in discovering new regulatory interactions not existing in the regulons. NetAct was applied using the regulons at different perturbation levels (25%, 50%, and 75%).

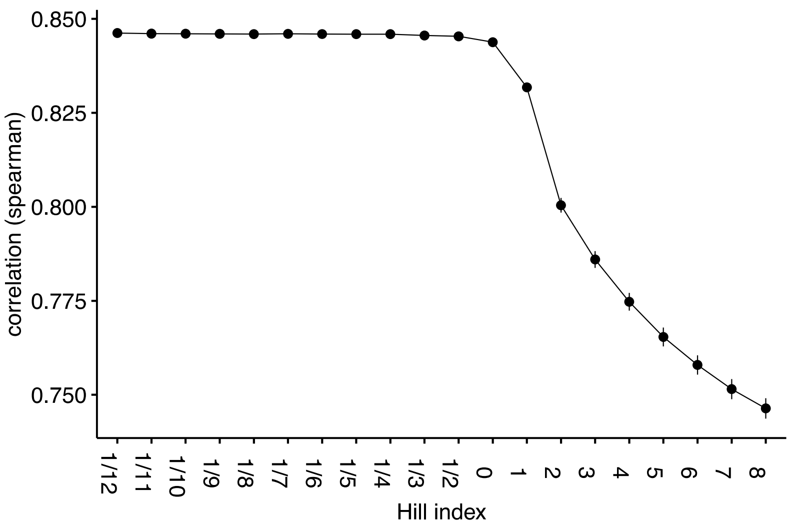


**Fig. S8. Optimization of the Hill coefficient in the TF activity inference.** We used in-silico benchmark dataset together with the synthetic TF regulatory datasets to optimize the Hill coefficient $n$ in the Hill function for inferring the TF activity. From a series of different Hill coefficients, we measured the performance by reporting the spearman correlation between the inferred activities and ground-truth activities. We found that when $n$ is small enough ($< \frac{1}{4}$), the performance remains comparably good.

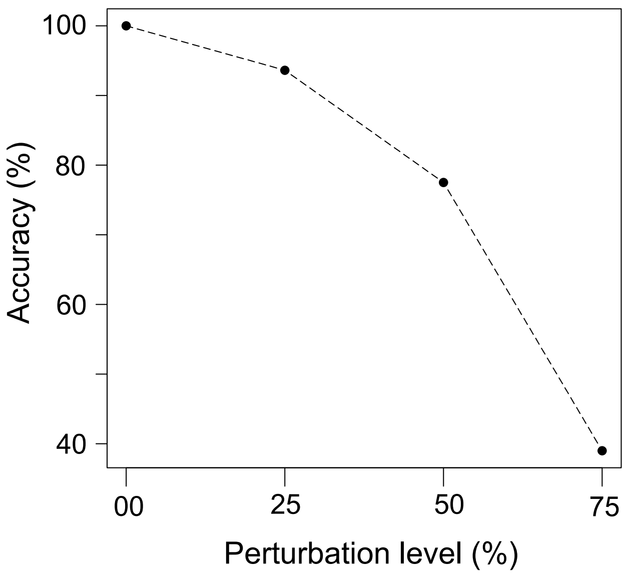


**Fig. S9. Comparison of NetAct grouping scheme for target genes with the synthetic gene regulatory network.** In this test, simulated gene expressions and regulons at different perturbation levels were provided to NetAct to find the grouping scheme for the targets of each TF. For targets in the ground-truth network, we evaluated the accuracy of the grouping scheme, *i.e.*, whether NetAct correctly predicts the activation/inhibition of the targets by the TF. Y-axis shows the accuracy defined as the fraction of TF-target pairs from NetAct whose activation/inhibition nature is consistent with the ground-truth network. X-axis shows the perturbation levels of the regulons used to calculate TF activities by NetAct.

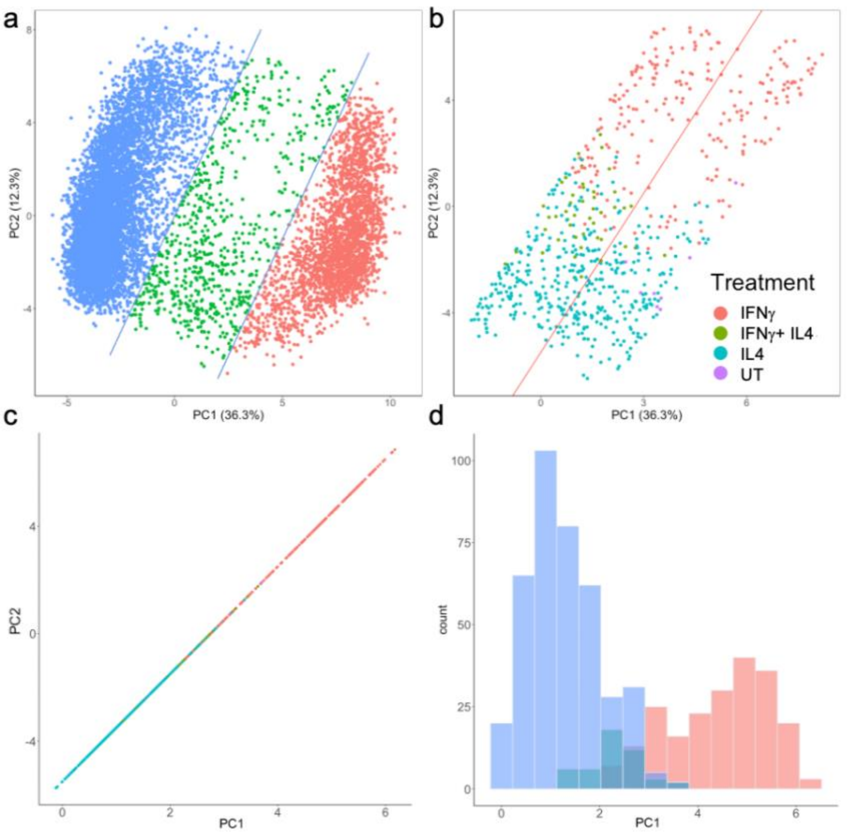


**Fig. S10. Analysis of 10,000 RACIPE-simulated gene expression profiles for the macrophage depolarization TF regulatory network**. **(a)** PCA projection of the simulated gene expression data from the macrophage network with lines (in blue) added to identify transitory models. **(b)** Transitory models identified in (a) with annotations of the experimental states; in addition, an average line of the two blue lines in (a) is added (in red). **(c)** orthogonal projection of all the points in (b) to the line in (b). **(d)** histogram of points along the line in (c). Together, this data shows that there are two main transition paths (IFNγ and IL4) and in between there are a lower density of models. (d) also shows that transitory models assigned to the hybrid state IFNγ_IL4 prefer a path more similar to the transitory models assigned to IL4.
